# origamiFISH allows universal, label-free, single molecule visualization of DNA origami nanodevices across biological samples

**DOI:** 10.1101/2022.09.19.508533

**Authors:** Wendy Xueyi Wang, Travis R. Douglas, Haiwang Zhang, Afrin Bhattacharya, Meghan Rothenbroker, Zhengping Jia, Julien Muffat, Yun Li, Leo Y. T. Chou

## Abstract

Structural DNA nanotechnology enables user-prescribed design of DNA nanostructures (DNs) for biological applications, but how DN design determines their bio-distribution and cellular interactions remain poorly understood. One challenge is that current methods for tracking DN fates *in situ*, including fluorescent-dye labeling, suffer from low sensitivity and dye-induced artifacts. Here we present origamiFISH, a label-free and universal method for single-molecule fluorescence detection of DNA origami nanostructures in cells and tissues. origamiFISH targets pan-DN scaffold sequences with hybridization chain reaction (HCR) probes to achieve thousand-fold signal amplification. We identify cell-type and shape-specific spatiotemporal uptake patterns within 1 minute of uptake and at picomolar DN concentrations, 10,000x lower than field standards. We additionally optimized compatibility with immunofluorescence and tissue clearing to visualize DN distribution within tissue cryo/vibratome-sections, slice cultures, and whole-mount organoids. Together, origamiFISH enables faithful mapping of DN interactions across subcellular and tissue barriers for guiding the development of DN-based therapeutics.

---

DNA origami nanostructures (DNs) enable rapid prototyping of highly monodisperse and user-prescribed nanoscale devices (Dey et al., 2021; Hong et al., 2017; Rothemund, 2006; Tseng et al., 2022). Combining compatibility for macromolecule decorations (i.e. proteins, antibodies, nucleic acids, small molecules etc.) with nanometer-scale precision of spatial arrangements and densities, DNs represent an attractive model for use in diverse applications such as drug delivery (Douglas et al., 2012; Li et al., 2018; Zhang et al., 2014), vaccines (Liu et al., 2021; Veneziano et al., 2020), biosensors (Han et al., 2019; Huang et al., 2018; Kosuri et al., 2019; Kuzuya et al., 2011, 2014), or as spatial organizers for biomolecules (Berger et al., 2021; Dong et al., 2021; Hawkes et al., 2019; Hellmeier et al., 2021; Kern et al., 2021; Veneziano et al., 2020). Many of these uses require controlling the cell uptake or subcellular localization patterns of DNs, such as cell-type and organelle-specific drug delivery or cell-surface display for receptor organization. Notably however, no existing imaging method allows for in situ detection of native, unmodified DNs. Field standards for DN imaging consist of transmission electron microscopy (TEM) and atomic force microscopy (AFM) for structural imaging (Bertosin et al., 2021; Endo and Sugiyama, 2018; Helmig et al., 2010; Kabiri et al., 2019; Silvester et al., 2021), or in vivo optical imaging for gross organ- and animal-level assessments (Ponnuswamy et al., 2017; Zhang et al., 2014). Alternatively, fluorescence imaging of DNs can be achieved through fluorophore-tagging using short, complementary strands of DNA, which has been useful for assessing the temporal distribution profiles of DN uptake within cells (Bastings et al., 2018; Liang et al., 2014; Wang et al., 2018). However, dye-labeling of DNs is challenged by a number of practical and design considerations: 1) mounting evidence suggests that fluorescence signals arising from DN-tagged dyes do not faithfully reflect the presence of intact DNs, and may instead arise from dissociated dyes or degradation products (Green et al., 2020; Koga et al., 2022; Lacroix et al., 2019); 2) fluorescent dyes have been shown to interact with biomolecules, cells and tissues, leading to altered cell uptake or biodistribution of tagged species (Hedegaard et al., 2018; Hughes et al., 2014; Koga et al., 2022; Zacharias et al., 2002); 3) fluorescent dyes are prone to sequence, pH or ROS-dependent quenching (Chen et al., 2008; Kretschy et al., 2016; Srisantitham et al., 2018; Zheng et al., 2014); 4) dye-tagging of DNs requires design and validations on a structure-by-structure basis, which is time and cost-limiting, especially for large-scale assays; and 5) imaging of dye-labeled DNs is constrained by the number of fluorescent tags introduced, thus limiting sensitivity for *in vivo* applications. Taken together, a practical method to faithfully visualize non-labeled DNs will enable new insights for the design and validation of DNA-based nanodevices for biological applications.

Here we introduce origamiFISH, a method for universal and sensitive detection of DNA origami nanostructures in cells and in tissues (Fig. 1). Through targeted imaging of conserved regions of the M13mp18 bacteriophage-based scaffold sequence, which appears in the vast majority of existing DNA origami designs (Dey et al., 2021; Rothemund, 2006), origamiFISH allows for universal detection of DNs without the need for structure-specific modifications (**Fig. 1a-d**). Additionally, by incorporating an in situ amplification step through hybridization chain reaction (Choi et al., 2014, 2018), origamiFISH allows for sensitive detection of single nanodevices across a variety of microscopy modalities (i.e. epifluorescence, slide scanner, confocal, and light-sheet microscopy), removing the entry barrier for DN imaging. We first demonstrate that origamiFISH has the sensitivity to detect DN uptake by cells under picomolar concentrations. Second, we show that origamiFISH performs robustly across a variety of DN shapes and cell-lines without additional parameter modifications or optimizations, revealing cell-type and structure-specific uptake patterns. Third, we chart the trajectory of DN uptake to identify shape-specific dynamics within the first minute of uptake. Fourth, we demonstrate that origamiFISH is compatible with immunohistochemistry for colocalization of DNs with cell-type and organelle-specific marker combinations. Finally, we extend origamiFISH to various tissue models, including 2D slice cultures of the mouse brain, cryosections of mouse lymph nodes, and 3D whole-mounts of human brain organoids. In summary, by decoupling DN imaging sensitivity from their degree-of-labeling, origamiFISH enables deeper and faithful profiling of DN biodistribution, which we envision to be useful for understanding their design rules and translational characterization for biomedical applications.

**Fig. 1.**
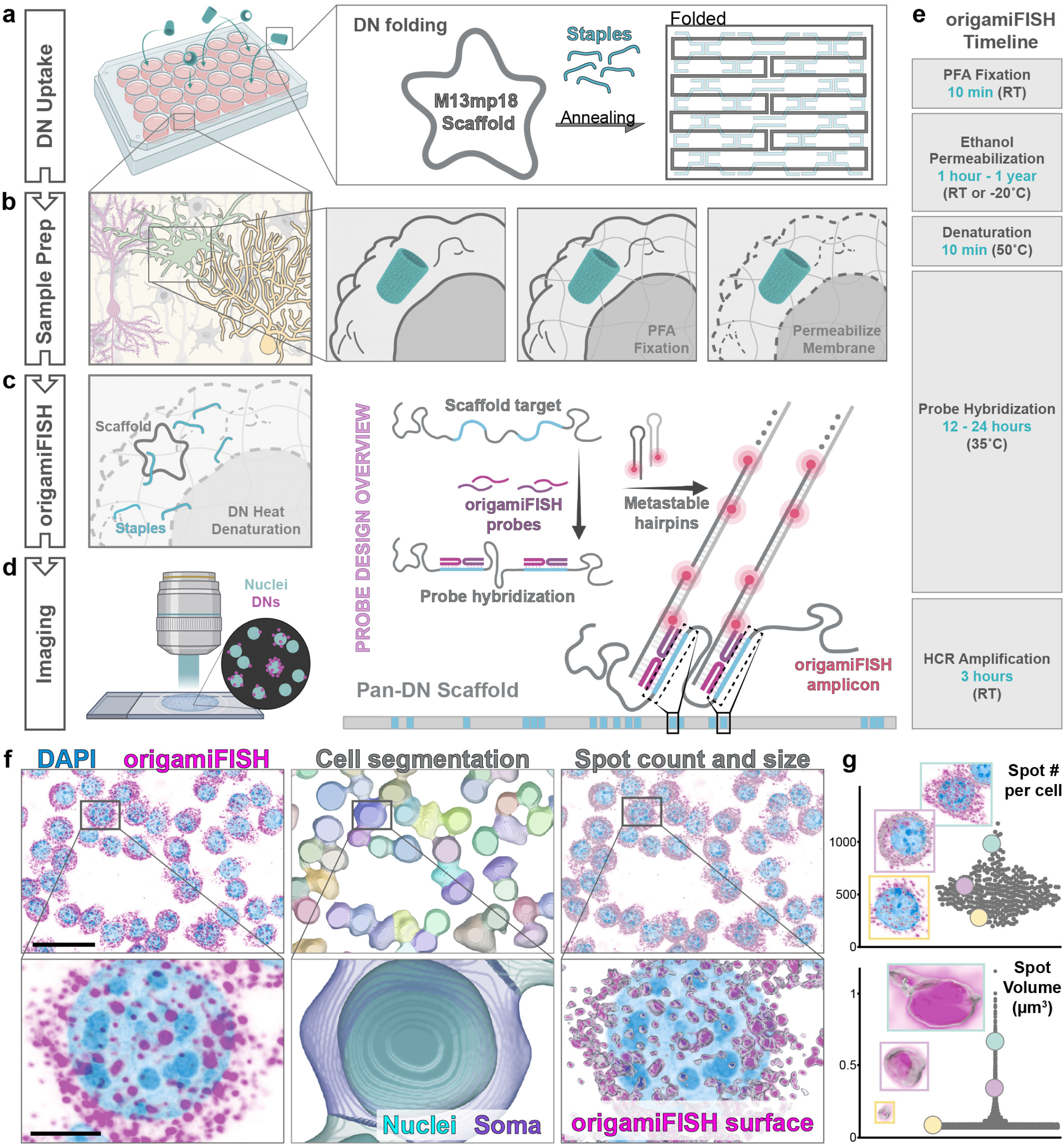
Overview of origamiFISH protocol and analyses. **a** DNs are introduced to cells, tissue slices, or organoids in 24- or 6-well plates. DNs are folded following standard DNA origami folding protocols, whereby the M13mp18-derived scaffold and structure-specific staples strands are thermo-annealed in the presence of MgCl_2_. **b** Samples comprising either cells on coverslips, tissue slices, or whole-mounts are subjected to the following procedures for sample preparation: samples were fixed in 4% paraformaldehyde for crosslinking of DNs and endogenous macromolecules in their in situ locations, followed by membrane permeabilization in 70% ethanol to enable probe entry. **c** DNs were denatured to dissociate scaffold and staple strands under heat and formamide conditions. origamiFISH probes were subsequently applied overnight to allow binding to the universal M13mp18-based scaffold. Hybridization chain reaction (HCR) hairpins were introduced next to for signal amplification. **d** Fluorescence imaging of origamiFISH stained samples with single cell, single molecule resolution. **e** overview of the origamiFISH protocol timeline. In total, origamiFISH can be completed in 1 day with between 1-2 hours of hands-on time. **f** Data analysis overview. Nuclei and cell segmentation is performed in Bitplane Imaris, followed by spot and surface reconstructions to allow single cell measurements of origamiFISH signal. Scale bars are 30 µm in the top panels and 5 µm in the bottom panels. **g** Dot plots of example data distribution in RAW264.7 cells following 30 minute DN barrel uptake. Example images demonstrate the variability in both DN spots per cell and volume of spots, which are robustly read out using origamiFISH.

## Development and validation of origamiFISH imaging

origamiFISH takes advantage of the presence of a long, circular scaffold DNA sequence that is present within DNA origami nanostructures (**Fig. 1a**) (Seeman and Sleiman, 2017; Tseng et al., 2022). In nearly all cases, DN scaffolds are derived from the 7249-nt genome of M13mp18 bacteriophage, with variants containing either sequence insertions (e.g. p7308, p7560, p8064, p8634) or deletions (e.g. p3024). To allow for structure-agnostic targeting of DNs, we designed origamiFISH probes by tiling along the conserved portion of the M13mp18 genome (**Fig. 1b-d; Supplementary Fig. 1**). Probes were designed following published guidelines for RNA single molecule fluorescence *in situ* hybridization (smFISH) (Eng et al., 2019; Raj et al., 2008). The workflow for origamiFISH is depicted in **Fig. 1**. As in RNA smFISH, we first ensured sample integrity through paraformaldehyde (PFA) fixation to crosslink endogenous and exogenous biomolecules to their in situ environments, and ethanol-based membrane permeabilization to allow probe entry (**Fig. 1b**). To expose intact double stranded DNs for binding by origamiFISH probes, we implemented a heat and formamide denaturation step, whereby short ssDNA staple strands are released from their scaffold counterparts (**Fig. 1c**). To enable sensitive detection of single DNs, we further utilized hybridization chain reaction (HCR) signal amplification chemistry, as has been previously demonstrated for single-molecule RNA detection (Choi et al., 2014, 2018). HCR allows for isothermal and enzyme-free amplification of fluorescent DNA polymers, following recognition of target-dependent initiator sequences (**Fig. 1c, d**). By tethering a HCR initiator to each DN-targeting probe, origamiFISH can theoretically achieve the equivalent of >1000 fluorophores per DN, making it feasible to detect single DNs in biological samples using conventional fluorescent microscopes (Choi et al., 2014, 2018; Tsuneoka and Funato, 2020). The complete origamiFISH protocol can be completed within 1 day, with ∼1.5 hours of hands-on time (**Fig. 1e**).

We first implemented origamiFISH in RAW264.7 macrophage cells, a widely utilized cell line within the DNA origami field with high DN uptake capacity (Kern et al., 2021; Liu et al., 2020; Maezawa et al., 2020; Zeng et al., 2022). Following our origamiFISH protocol, DNs appear as bright, distinct fluorescent puncta, or “spots”, within each cell (**Fig. 1f**). To analyze these spots and quantify the uptake pattern of DNs, we established an image analysis pipeline within the Bitplane Imaris software which performs semi-automated cell segmentation, spot detection, and spot quantifications (**Fig. 1f**). This allowed us to extract quantitative features of DN uptake with single-cell resolution, including the number of DN spots per cell, size and intensity of spots, and differential DN localization within the cytoplasm versus nucleus (**Fig. 1g**). Initial origamiFISH experiments revealed a high degree of variability in DN uptake at the single cell and single molecule level even within the same sample (**Fig. 1g**).

To optimize origamiFISH, we first tested two probe design strategies (**Fig. 2a**). We designed 25 single stranded DNA probes (HCR v2.0) which span a total of 625nt (Choi et al., 2014), and 20 split-initiator probes (HCR v3.0) which span a total of 1000nt along the M13mp18-based scaffold sequence (**Supplementary Fig. 1**) (Choi et al., 2018). Although both single and split-initiator probes produced a robust signal, we observed that origamiFISH background fluorescence was visibly reduced within negative no-DN controls, when probed using split-initiator probes (**Fig. 2a**), consistent with published findings for RNA targets (Choi et al., 2018). We therefore proceeded with using the split-initiator origamiFISH probe set for the remainder of this study, which was additionally confirmed to be reliable across a variety of imaging modalities (**Supplementary Fig. 2**).

**Fig. 2.**
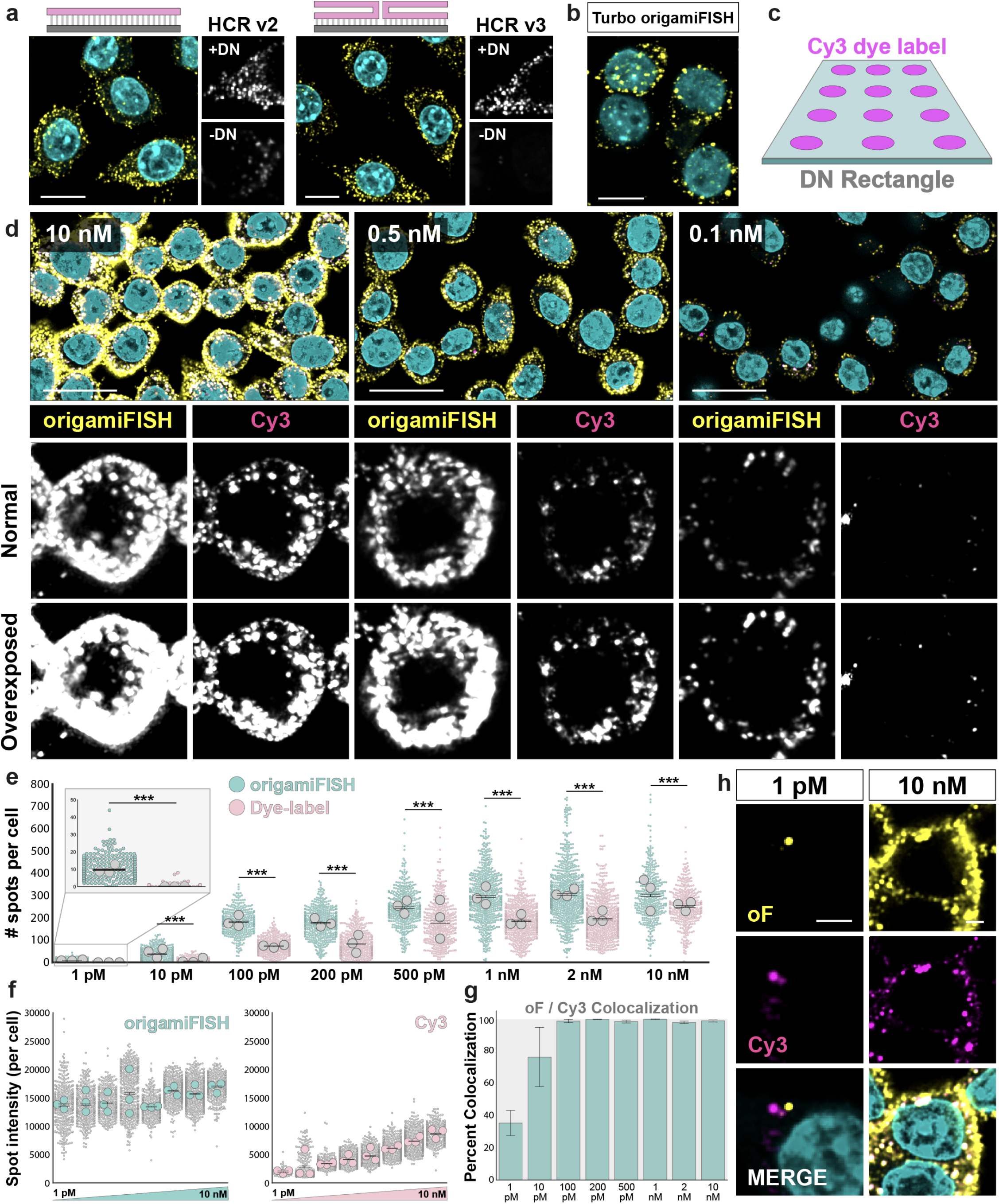
origamiFISH enables highly sensitive imaging of DNA origami nanostructures in situ. **a** Comparison of HCR version 2 single stranded DNA (ssDNA) probes and HCR version 3 split initiator probes (Choi et al., 2014, 2018). A schematic of the general probe design strategy is shown, with the DN scaffold target in gray and complementary origamiFISH probes in pink. Although both HCR v2 and v3 probes produce a robust signal, split initiator probes significantly diminish background staining in negative no-DN control samples. **b** Representative image of turbo origamiFISH. Image shows RAW264.7 cells following 30 minute DN rectangle uptake in (**a, b**). **c** Schematic of Cy3+ DN rectangle. 12 Cy3 dyes are tethered to each DN rectangle on one side through complementary DNA handle/anti-handle chemistry. **d** Images of RAW264.7 cells following 30 minute uptake of Cy3+ DNA origami rectangle at various concentrations. *Top*: representative merged images with DAPI (cyan), origamiFISH (yellow), and Cy3 (magenta). *Bottom*: zoomed in views of single cells from the top panel. **e** Dot plot showing quantifications of single cell origamiFISH (teal) and Cy3 (pink) spot counts across concentration gradients. Each colored dot represents quantifications from a single cell, and each gray dot represents cumulative average for each biological replicate. origamiFISH resulted in significantly increased spot counts for all conditions. ***p < 0.001. Means of two groups were compared using the Mann-Whitney two-tailed non-parametric test. **f** Dot plot quantifications for origamiFISH and Cy3 signal across concentrations (16 bit image). Gray dots represent average spot intensity per cell. Colored dots represent average intensity for each biological replicate. **g** Percent colocalization rates between origamiFISH+ and Cy3+ spots across concentrations. Positive colocalization (cyan bars) represent Cy3+ spots which are additionally positive for origamiFISH. Negative colocalization (gray negative space) represent Cy3+ spots which are negative for origamiFISH. **h** Representative images at 1pM and 10nM, respectively. Examples of both Cy3+/origamiFISH- and origamiFISH+/Cy3-spots are observed at 1pM, but not 10nM. Scale bars are 10 µm in all panels except (**h**), which is 1 µm.

It has previously been demonstrated that smFISH protocols can be shortened from one day to 15 minutes using a combination of methanol sample fixation and ultra-high probe concentrations (Shaffer et al., 2013). To test whether origamiFISH can be similarly accelerated, we established protocols for Turbo-origamiFISH, which enables end-to-end origamiFISH in 1 hour (**Fig. 2b; Supplementary Fig. 3**). Turbo-origamiFISH produced bright and distinct puncta in cells which are dependent on methanol fixation, and displayed similar subcellular localization patterns compared to origamiFISH (**Supplementary Fig. 3**). We note that Turbo-origamiFISH results in visibly reduced puncta numbers compared to origamiFISH, suggesting sacrificed detection sensitivity. However, we envision that Turbo-origamiFISH will offer unique advantages for rapid and large-scale prototyping of DNs for cellular applications which may be difficult with more time-consuming protocols. Together, origamiFISH enables robust, quantitative and high-quality imaging of DN biodistribution with single cell and single molecule resolution in situ.

### Sensitive detection of picomolar ranges in DN concentrations

With the optimized protocol, we next investigated the detection sensitivity of origamiFISH. Typical DN concentrations used for cellular uptake assays range between 10nM to 250nM (Lacroix et al., 2019; Ponnuswamy et al., 2017; Rajwar et al., 2022; Wang et al., 2018), whereas single-molecule sensitivity is well-established for smFISH (Raj et al., 2008). To determine if these high uptake concentrations were necessitated by the limited brightness of standard dye labeling approaches, we sought to compare signals arising from origamiFISH and Cy3 dye labels within the same sample. To this end, we designed a Cy3-labeled DN rectangle, decorated with 12 Cy3 dyes on one side (**Fig. 2c**). We tested a spectrum of Cy3+ DN concentrations in RAW264.7 cells, ranging from 1 picomolar to 10 nanomolar (**Fig. 2d; Supplementary Fig. 6a**). Prior to the cell experiments, DNs were coated with the polymer oligolysine-poly(ethylene glycol) (1:1 N/P ratio; hereafter K_10_-PEG) as previously demonstrated, to prevent digestion by nucleases as a confounding factor (**Supplementary Fig. 4, 5**) (Ponnuswamy et al., 2017). Both origamiFISH and Cy3 dyes generally recapitulated the gradient of concentrations as expected, with the number of DN spots increasing with DN concentration (**Fig. 2e, inset**). However, origamiFISH detected significantly more DN spots per cell compared to Cy3, across all concentrations analyzed (**Fig. 2d, e; Supplementary Fig. 6a, b**). This is particularly striking at lower uptake concentrations, where origamiFISH detected a mean of 9.76 (± 0.29 s.e.m) DN spots per cell following 1pM uptake, compared to a mean of 0.15 (± 0.03) spots detected by Cy3 (**Fig. 2e**). Additionally, the mean spot intensity for origamiFISH with 1pM DN uptake was 2.7 times higher compared to Cy3 intensity in the 10nM uptake condition (**Fig. 2f**; 1pM origamiFISH: 13714 ± 186; 10nM Cy3: 5070 ± 243; s.e.m; 16-bit image). Both origamiFISH and Cy3 signals demonstrate saturation past 2nM DN uptake (**Supplementary Fig. 6c**). Although we cannot directly correlate signal intensity of the two imaging approaches due to inherent differences in dye characteristics, these results strongly suggest that origamiFISH is orders of magnitude more sensitive compared to traditional dye-labeling strategies. We note that our experiments additionally incorporated a shorter-than-recommended HCR amplification time of three hours (Choi et al., 2014, 2018). Consequently, our results represent the lower detection range of what is technically possible with the HCR approach. We maintained a short amplification time to limit spot sizes for optimized image analyses, and to shorten the experimental timeline. However, origamiFISH signals can be further enhanced by increasing HCR hairpin concentration or amplification time.

We next analyzed the colocalization between origamiFISH and Cy3 signals as a function of DN uptake concentration. We observed near 100% colocalization at DN concentrations ranging from 10 nM down to 100 pM, but a striking loss of colocalization at DN concentrations below 100 pM (**Fig. 2g, h**). Following 1pM uptake, we noted many instances of both origamiFISH+/Cy3- and Cy3+/origamiFISH-spots (**Fig. 2h; Supplementary Fig. 7**). We reasoned that although the lack of Cy3 signal in origamiFISH+ spots could be due to differences in fluorescence detection sensitivity of the two methods, the presence of Cy3+ spots without origamiFISH signal was more perplexing, suggesting the dissociation of DN scaffold (targeted by origamiFISH) and Cy3+ staples strands. To assay this possibility, we differentially quantified the proportion of Cy3+ spots which were positive or negative for origamiFISH. Interestingly, we found that the majority of Cy3+ spots were negative for origamiFISH signals following 1pM uptake (**Fig. 2g**). These results suggest that, under these dilute conditions, either DN degradation of Cy3 staple dissociation, or both, is detectable within the first 30 minutes of cell uptake, despite protective K_10_-PEG coatings. We cannot exclude the possibility of DN degradation at higher concentrations as the rescue in colocalization rates may be biased by signal accumulation within endosomal compartments. Nevertheless, these results have implications for the stability of DNs under dilute concentrations, such as following their *in vivo* administration and distribution to target tissues of interest. Our findings caution careful study design and characterizations of DN stability for *in vitro* and *in vivo* applications. It has been proposed that strategies such as glutaraldehyde cross-linking of oligolysine coatings can additionally increase DN stability ∼250 fold under high DNase concentrations (Anastassacos et al., 2020). Similarly, UV-induced point welding of DNA strands within DNs has also been developed (Gerling et al., 2018). By demonstrating the utility of origamiFISH in detecting DN integrity under cellular conditions, we envision it will complement other assays, such as those based on Forster resonance energy transfer (FRET) (Ponnuswamy et al., 2017) or TEM (Wang et al., 2018), in evaluating various DN stabilization strategies under biological environments.

### Profiling of cell-type and shape specific DN uptake patterns

We next asked whether origamiFISH can be used to interrogate DN-specific uptake patterns across different shapes (rectangle, barrel, and long rod), and cell lines (RAW264.7 macrophages and HEK293 embryonic kidney cells). We selected shapes spanning one or more dimensions, including representative 1D (long rod), 2D (rectangle) and 3D (barrel) DNs, to identify design parameters for informing future studies of DN biology (**Fig. 3a**). DNs were synthesized and characterized by TEM and agarose gel electrophoresis (**Supplementary Fig. 4, 5**). We performed DN uptake in cell culture media containing 1nM DN concentrations to ensure sufficiently quantifiable signals under low uptake conditions. RAW264.7 cells displayed significantly enhanced uptake efficiency across DNs compared to HEK293 cells, consistent with their role as phagocytes (**Fig. 3b, c**). Additionally, we observed DN shape-specific rules for cellular uptake. Although the theoretical surface area for DN rectangles (60 × 90nm; ∼10800nm2) and barrels (30 × 60nm; ∼10900nm2) are comparable, we observed significantly increased levels of DN uptake for rectangles in RAW264.7 cells. This trend was reversed in HEK293 cells, where DN barrels displayed a slight, but significant increase in uptake efficiency. By contrast, uptake of DN long rods was much decreased in both RAW264.7 and HEK293 cells (**Fig. 3c**). As long rods are 400 nm in length, this is likely due to increased energy costs to invaginate and engulf longer DNs with higher aspect ratios (Bastings et al., 2018; Champion and Mitragotri, 2006). Together, these results demonstrate the use of origamiFISH for enabling label-free DN imaging across cell-lines and DN shapes without additional optimizations. origamiFISH identifies both shape and cell-type specific uptake dynamics. Variability in uptake trends between cell-types additionally suggest that DNs may likely engage cell-type dependent uptake mechanisms and pathways.

**Fig. 3.**
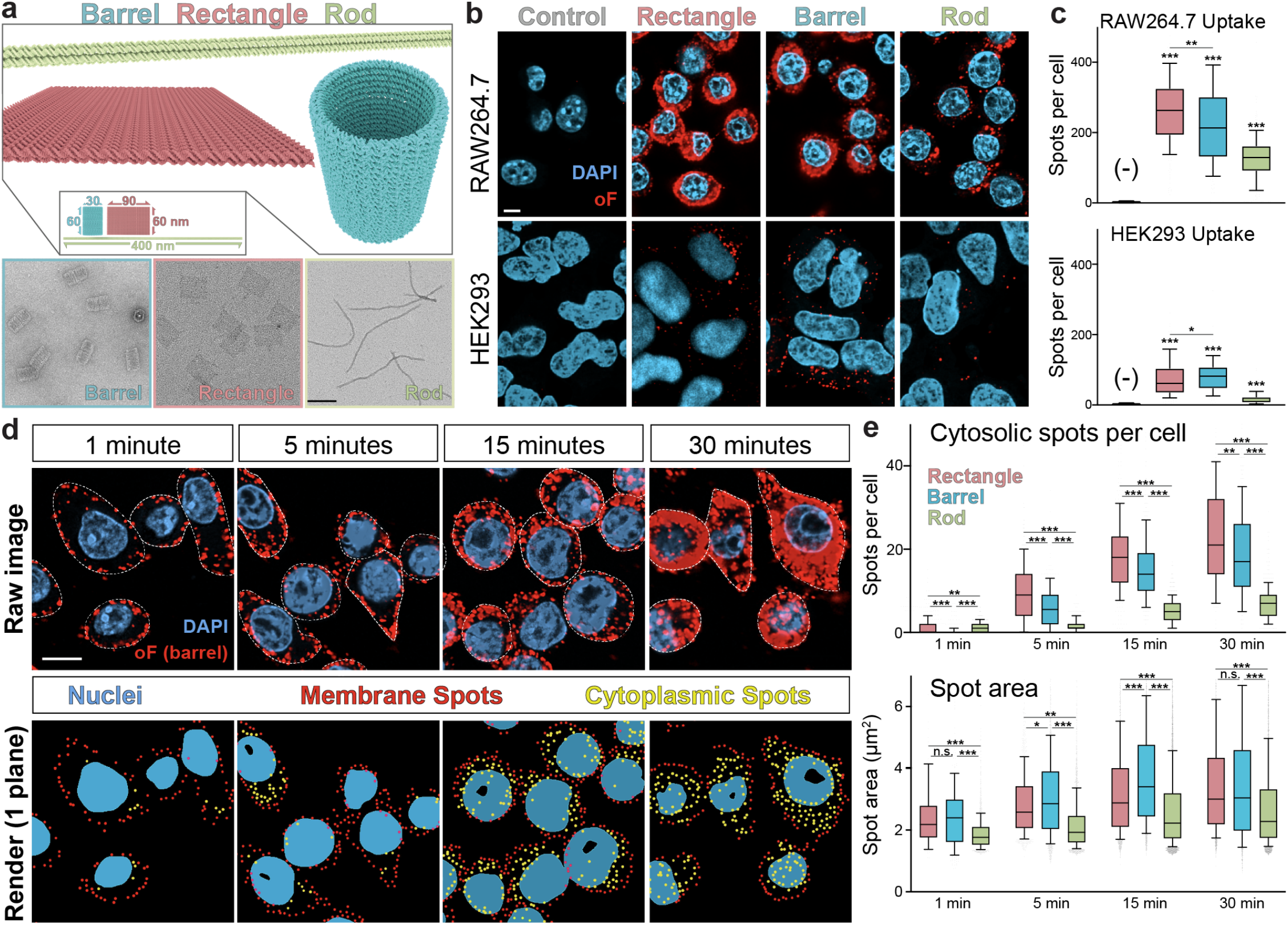
origamiFISH reveals cell-type, shape, and time-dependent uptake patterns of DNs. **a** Schematic of DN shapes used in this study. DN barrel (teal; 60 × 30nm), rectangle (pink; 60 × 90nm) and rod (green; 400 × 7nm) are folded following standard DNA origami thermo-annealing protocols as outlined in the methods section. Transmission electron micrographs of the respective shapes are shown in the bottom panel. Scale bar is 50nm. **b** Representative origamiFISH images following DN uptake of three shapes and two cell lines. **c** Box plot quantifications for the number of origamiFISH spots across shapes (rectangle: pink bars; Barrel: teal bars; Long rod: green bars) and cell lines. Negative no-DN control quantifications are shown in the first bar (-). Lines in box plots denote 25th percentile, median and 75th percentile respectively. Whiskers represent 90th and 10th percentile. **d** Dynamics of the first 30 minutes of DN uptake. Shown are representative images of barrel uptake. *Top*: Raw images for the respective timepoints. Cell boundaries as determined by cellular autofluorescence are outlined with dashed white lines. *Bottom*: Render of spot localizations, with membrane localized spots along the boundary of cell membrane shown in red and cytosolically localized spots shown in yellow. Raw images are maximum projections of 5 z-planes, while renders are created using 1 z-plane to de-crowd dense regions. Scale bars are 5µm in (**b, d**). **e** Box plot quantifications for the number of spots per cell (*top*) and spot area (*bottom*) across shapes and uptake times. *p<0.05, **p<0.005, ***p<0.00001. Means of two groups were compared using the Mann-Whitney two-tailed non-parametric test.

### origamiFISH reveals DN shape-specific uptake trajectories within 1 minute of uptake

Cellular engulfment and endocytosis of foreign nanoparticles occur on the order of minutes (Mager et al., 2012), but past studies of DN uptake have mostly focused on longer-term assays, over the course of hours or days (Bastings et al., 2018; Ponnuswamy et al., 2017). One study which visualized DN uptake over 4 days found negligible uptake 5 minutes post DN introduction, despite a 250 nM concentration (Wang et al., 2018). We wondered if the lack of short time-scale assays using dye-labeled DNs may be due to requirements for signal accumulation within endocytic compartments, and if origamiFISH may offer the sensitivity necessary for probing the early phase of DN uptake biology. To this end, we performed time-course assays for all three DN shapes, over the course of 30 minutes (1, 5, 15, and 30 minutes). In this set of experiments, we addtionally introduced DNs at 0.2nM concentration to highlight the detection sensitivity of origamiFISH.

origamiFISH detected ample accumulation of DN signal following just 1 minute of uptake (**Fig. 3d**). The number of origamiFISH+ DN spots increased with uptake time across all shapes analyzed as expected (**Fig. 3e; Supplementary Fig. 8**). Additionally, DN spots displayed corresponding increases in size, suggesting accumulation within endosomal or lysosomal compartments. Intriguingly, we noted that origamiFISH signal appeared to be largely membrane-localized at early time points (i.e. 1 and 5 minute), followed by gradually increasing numbers of internalized, cytosol-localized spots as uptake progressed (**Fig. 3d, e**). To obtain a quantitative readout for these spatiotemporal trends, we implemented an image analysis pipeline to differentially quantify the number of membrane- and cytosol-localized spots at the single cell level (**Supplementary Fig. 8a**; see methods). Number of membrane-localized spots were obtained by proxy from quantifications of the number of origamiFISH spots which localized to cell boundaries, as segmented by cellular autofluorescence. We observed highly dynamic, shape-specific differences in DN internalization kinetics as soon as 1 minute following uptake (**Fig. 3e**). Importantly, these early differences in shape-specific uptake did not match perfectly with trends observed at later time points. For instance, DN barrels displayed the lowest rate of uptake within the first minute (**Fig. 3e; Supplementary Fig. 8b**). Specifically, we observed a mean of 1.32 (± 0.15 s.e.m) cytosolic origamiFISH spots for DN rectangles, compared to a mean of 0.37 (± 0.06) for barrels and 1.08 (± 0.05) for long rods (**Fig. 3e**). This result suggests that barrel internalization may be rate-limiting within the first minute of DN introduction. However, between 1 to 5 minutes, the number of internalized barrel spots displayed a 16.7 fold increase to 6.24 ± 0.30 spots per cell, compared with 7.7 fold (10.13 ± 0.46 spots at 5 min) for rectangles and 1.57 fold (1.69 ± 0.06 spots) for long rods. Increases in cytosolic spots appear to be largely independent of membrane localization, as the number of membrane-localized spots are relatively stable throughout the full 30 minute time course (**Supplementary Fig. 8c**). We additionally observed that membrane adsorption is structure-specific, with DN rectangles and barrels accumulating significant membrane staining within the first minute of uptake (**Supplementary Fig. 8b, c**). By contrast, we did not observe significant membrane-localized staining in DN long rod samples at any time point. These results provide a glimpse into a novel, early phase of DN uptake biology and shape-dependent intracellular trafficking, and provide corresponding tools for elucidating downstream mechanisms in future studies.

### origamiFISH is compatible with immunohistochemistry across 2D and 3D tissue models

Developing DNs for biological applications requires visualizing their uptake and distribution patterns within the context of the surrounding cellular and tissue environment. For instance, cancer or immune cell targeted therapeutics need to make their way towards their respective target cell-types. On a finer-scale, endosomal or membrane-targeted DNs additionally need to reach their respective subcellular compartments. Protein labeling using immunofluorescence is commonly used for labeling of cell-type and organelle-specific markers. Therefore, we next tested whether origamiFISH can be combined with antibody-assisted immunolabeling for colocalizing DNs with proteins across cell and tissue models (**Fig. 4a**).

**Fig. 4.**
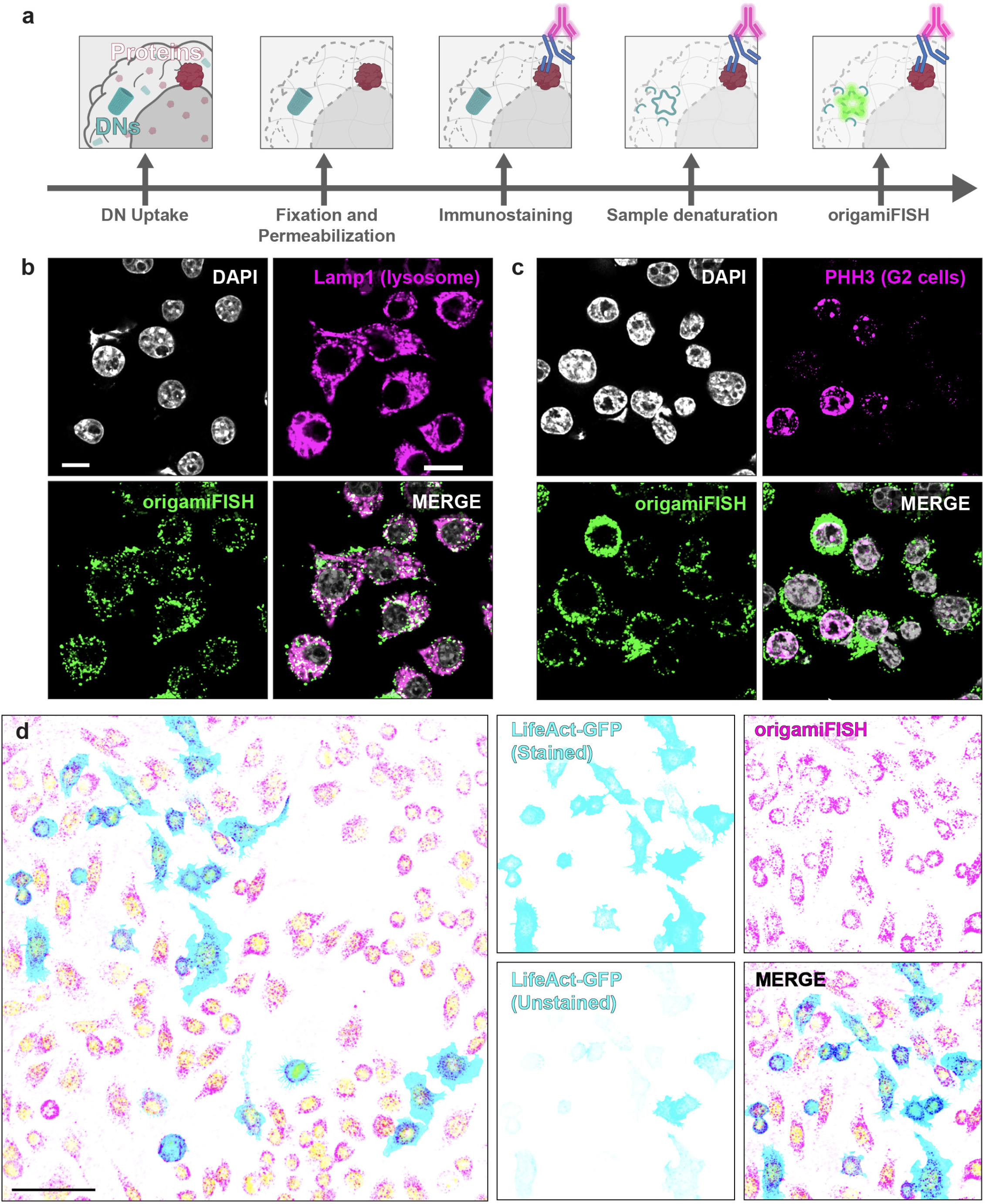
origamiFISH is compatible with immunohistochemistry. **a** Workflow overview for combining origamiFISH and protein immunostaining. Immunostaining using antibodies of choice is performed first, followed by washes and origamiFISH. **b** origamiFISH (green) and Lamp1 (magenta) co-stain in RAW264.7 cells following 30 minutes of DN rectangle uptake. High degree of colocalization is observed between origamiFISH and Lamp1+ lysosomes. **c** origamiFISH (green) and PHH3 (magenta) co-stained in RAW264.7 cells following 30 minutes of DN rectangle uptake. PHH3+ mitotic cells display visibly fewer origamiFISH spots. **d** origamiFISH (magenta) and GFP (cyan) co-stained in RAW264.7 cells transfected with LifeAct-GFP plasmids. Native GFP fluorescence is additionally compared with antibody-amplified GFP signal. Scale bars are 50µm.

It has previously been shown that the vast majority of nanoparticles localize to Lamp1+ lysosomes as their final destination within cells (Shapero et al., 2011). Consequently, targeting of endo-lysosomal receptors is emerging as an important strategy in nanoparticle-based therapeutics (Comberlato et al., 2022; Tseng et al., 2022). To visualize if DNs similarly traffic to lysosomal compartments, we co-stained origamiFISH with Lamp1 antibodies following 30 minute uptake of DN rectangles in RAW264.7 cells. Similar to silica and gold nanoparticles, we observed a high degree of colocalization between origamiFISH and Lamp1 puncta, confirming the lysosomal fate for DNs (**Fig. 4b**). We additionally co-stained similar samples for phosphohistone H3 (PHH3), a marker for mitotic cells in the late G2 and M phase of the cell cycle (Hendzel et al., 1997), to ask if cells in varying stages of division display differential DN levels. We observed that high PHH3 expression typically correlated with lower DN uptake, suggesting potential cell cycle dependent uptake mechanisms (**Fig. 4c**). Lastly, the time course study of DN uptake prompted us to ask if antibody-assisted amplification of fluorescent proteins could be combined with transfection of compartment-specific plasmids to improve cell and membrane segmentation for origamiFISH image analysis. We reasoned that a sparse cytosol or membrane marker would substantially improve accuracy of single cell segmentation. To this end, we transfected RAW264.7 cells with LifeAct-GFP plasmid for whole cell labeling prior to performing DN uptake (**Fig. 4d**). Although native GFP fluorescence diminished following origamiFISH, we demonstrate that amplification with GFP antibody enables transfection-based cell segmentation. Together, these results demonstrate the compatibility between origamiFISH and immunohistochemistry.

We next asked whether origamiFISH could be used to interrogate DN biology in tissue. To our knowledge, there are no existing studies on the characterization of cell-type specific DN uptake *in vivo*. One challenge is the poor sensitivity of dye labels as there is a limited number of dyes that can be tagged onto each DN (typically 10 to 20). This is additionally complicated by low DN concentrations in target tissues following their *in vivo* distribution. Furthermore, tagged dye species can interact nonspecifically with endogenous biomolecules, and dissociate from DNs *in vivo* (Hughes et al., 2014; Lacroix et al., 2019). We first extended DN uptake and origamiFISH to mouse draining lymph nodes, an immune organ important for sequestration of antigens following vaccination, and a key site of interest for nanovaccines (Liu et al., 2021; Zhang et al., 2020). We performed intramuscular injection of DN barrels in C57BL/6 mice, and harvested the inguinal lymph nodes (LNs) 4 hours post-injection. Following post-fixation, LNs were sectioned on a vibratome to prepare 100µm tissue sections. We performed origamiFISH on free-floating LN sections following our optimized protocol, with the following adaptations: 1) DN denaturation was performed at 60°C for 15 minutes to account for potential loss of formamide penetration; 2) HCR amplification was performed overnight to ensure sufficient signal amplification in tissue slices. origamiFISH revealed region and cell-type specific rules to DN uptake in LNs. We noted that cells with high DN uptake (hereafter origamiFISH^HIGH^) were distributed throughout the LN, but were highly enriched around B-cell follicles (**Fig. 5a**). Specifically, we observed that DNs were taken up by large numbers of cells around the follicle periphery, consistently displaying large soma morphologies (**Fig. 5a, inset i**). Fewer numbers of small origamiFISH^HIGH^ cells were additionally present within the B-cell follicles (**Fig. 5a, inset ii**). To determine if the large cells around the follicle periphery were macrophages, we co-stained LNs with F4/80 and origamiFISH (**Fig. 5b**). We confirmed that the majority of origamiFISH^HIGH^ cells were F4/80+ macrophages. However, we noted that only a subset of F4/80+ cells were additionally origamiFISH+, suggesting subtype-specific differences in DN uptake capacity within LN macrophages. Lastly, we noted examples of origamiFISH^HIGH^; F4/80-cells, suggesting that DNs are sequestered by multiple cell populations within LNs (**Fig. 5b**); future work will elucidate the diversity of these cellular repertoires. Together, these results demonstrate the distribution pattern of DN targeting to LNs, and extend dual origamiFISH and protein imaging to mouse tissue.

**Fig. 5.**
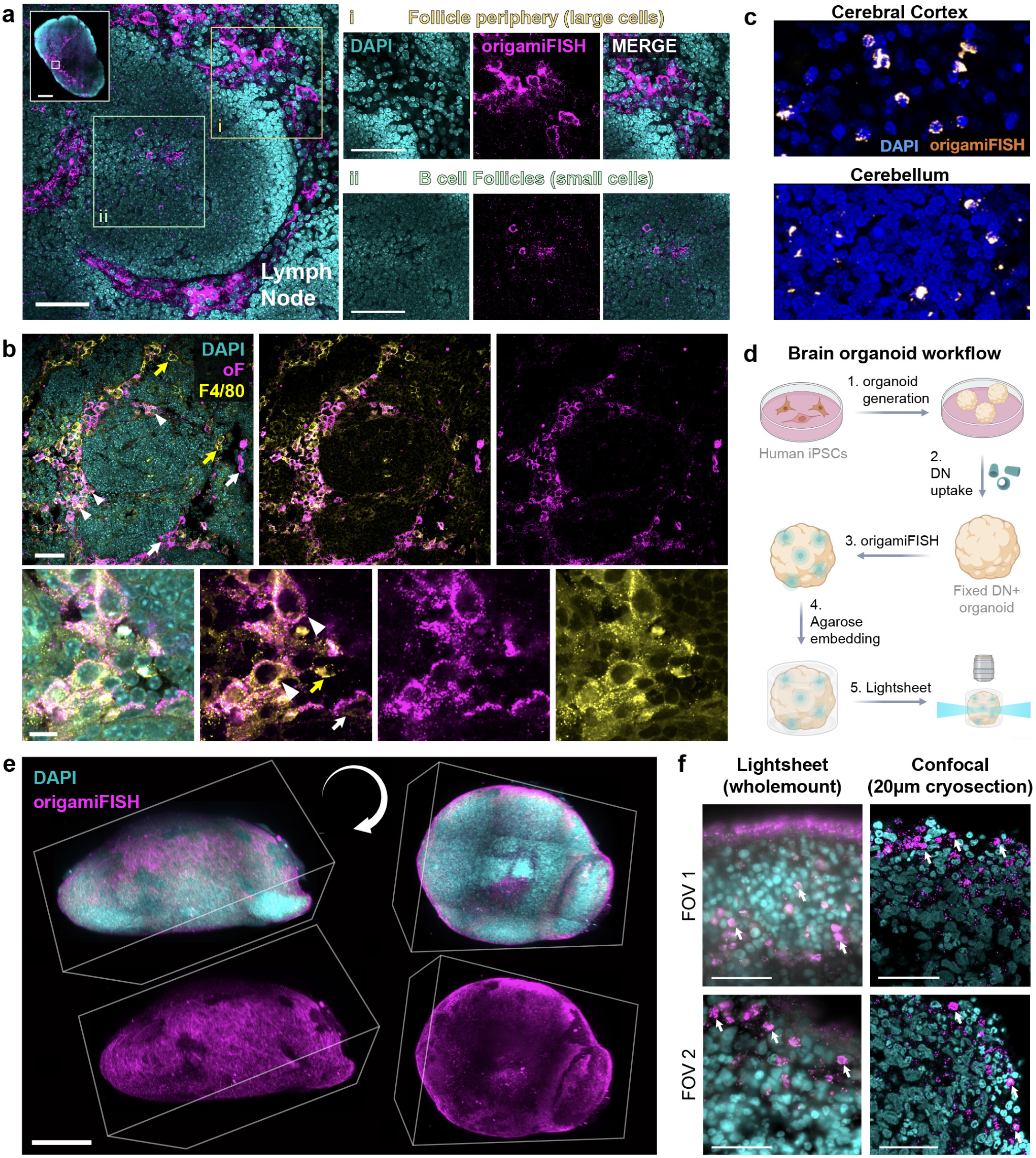
origamiFISH is robust across tissue models. **a** origamiFISH (magenta; DN barrels) labels subsets of cells in the mouse lymph node. *Left*: representative view of a B-cell follicle within the lymph node. origamiFISHHIGH cells with large soma morphology are enriched around the periphery of the B-cell follicle, and are additionally shown in the yellow inset (**i**). Smaller numbers of origamiFISHHIGH cells are additionally observed within B-cell follicles and are additionally shown in the green inset (**ii**). **b** Dual origamiFISH (magenta) and IHC identifies DNs in F4/80+ (yellow) macrophages. Yellow arrows denote F4/80+ macrophages which are negative for origamiFISH. White arrows denote origamiFISH+ cells which are F4/80-. White arrowheads denote cells double positive for origamiFISH and F4/80. **c** Representative images of the cerebral cortex (*top*) and cerebellum (*bottom*) following 15 minute DN barrel uptake. A subset of cells in both brain regions are high for origamiFISH (orange). **d** Workflow for whole-mount organoid DN uptake and imaging. Cerebral organoids are induced from human iPSC cells following established protocols. At around 12 weeks of age, DN barrels were introduced for 4 hours. origamiFISH was performed with integrated tissue-clearing in whole organoids (∼1-2mm in diameter). Agarose embedded organoids are imaged on a lightsheet microscope. **e** Representative lightsheet perspective views of whole cerebral organoids. origamiFISH (magenta), DAPI (cyan). Box represents image boundaries. Two perspective views are shown on the left and right of the same organoid. origamiFISH signal is enriched within the surface of the organoid. **f** Left: representative lighsheet image planes from (**e**). Cells with high DN uptake are denoted with white arrows. Right: Representative confocal image planes. Scale bars are 100 µm in (**a**), 500 µm in (**a**; whole LN inset), 100 µm in (**b**; top), 10 µm in (**b**; bottom), 50 µm in (**e, f**)

We next validated origamiFISH for the mouse brain and human brain organoids due to a few considerations. First, the brain is recognized as one of the most challenging tissue models for fluorescence imaging, due to its high fat content, tissue autofluorescence and extensive neurite networks (Schnell et al., 1999). We reasoned that although it was not feasible to validate origamiFISH across all tissue systems, approaching a challenging model as proof-of-concept may reveal areas which require additional protocol optimizations. Furthermore, previous studies have presented evidence for charge-selective nanoparticle uptake by neuronal populations (Dante et al., 2017). As inherently charged entities, DNA-based nanoparticles may offer a uniquely attractive toolbox for targeted delivery to the brain. As a first step towards this long-term goal, we aimed to validate origamiFISH in the nervous system. To this end, we introduced DN barrels to acutely prepared live slices of the mouse brain over the course of 15 minutes to probe uptake specificity in cells of the cerebral cortex and cerebellum (**Fig. 5c**). In both regions, we observed sparse but distinct DN uptake in small populations of cells, revealing DN uptake patterns within the nervous system. Future studies will examine the cell types responsible for DN uptake.

Finally, we applied origamiFISH towards whole-mount preparations of human brain organoids (**Fig. 5d**). Three-dimensional human iPSC-derived organoid models have emerged as a tractable model system for addressing diverse questions in neuro and cancer biology (Kim et al., 2020). We reasoned that organoids can similarly enable the design of high-throughput assays for validating therapeutic DNs in a humanized model. Additionally, we aimed to extend origamiFISH towards intact 3D tissues as another demonstration of its versatility, and to assay DN spatial distribution, accessibility, and penetration in non-vascularized tissue systems. Towards this end, we introduced DN barrels to 12 week old cortical brain organoids, a maturity stage containing diverse neurons and neural precursors, in addition to smaller numbers of astrocytes (Pa$ca et al., 2015). Following 4 hour uptake, organoids were postfixed prior to undergoing a modified 7-day origamiFISH protocol with integrated tissue clearing to enable whole organoid imaging on a lightsheet microscope (see methods). origamiFISH revealed spatial distribution patterns of DNs within the intact organoid (**Fig. 5e**). We observed high DN uptake by select cells enriched within the outermost 150µm of the organoid surface (**Fig. 5f**). To ensure that the restriction of DN uptake to cells within the organoid periphery was not due to the lack of probe penetration in whole-mount preparations, we confirmed these results using confocal imaging of organoid cryosections, which demonstrated similar limitations in DN penetration (**Fig. 5f**). These results corroborate previous observations of DN accessibility and penetration limits within cancer spheroids (Wang et al., 2021a), and prompt the need for more detailed characterizations of DN design rules for reaching deeper tissue regions within poorly vascularized organoid, spheroid, and tumor models. Taken together, our results demonstrate the utility of origamiFISH in capturing the faithful localization patterns of DNs within intact tissue systems, and offer a tool for characterizing DN design parameters for reaching deep tissue regions. Additionally, these results demonstrate the compatibility of origamiFISH to probe DN uptake across diverse tissue systems.

## Conclusions

In summary, we present origamiFISH as a novel imaging approach for label-free visualization of DNA origami nanostructures *in situ*. We demonstrate the utility and sensitivity of origamiFISH across multiple cell lines and tissue models in detecting: 1) cell uptake under picomolar concentrations of DNs; 2) differences in DN and fluorophore tag localization; 3) shape and cell-type specific rules in DN uptake; 4) early phases of DN uptake dynamics, as soon as 1 minute following uptake, and 5) DN profiling in whole-mount preparations. As with any technology, there are some limitations. First, performing origamiFISH requires sample fixation and therefore cannot be combined with live cell observations and tracking of DN uptake. Second, while origamiFISH more faithfully maps DN localization in biological tissues compared to dye-labeling, and provides evidence for DN degradation under picomolar concentrations, discriminating between fully and partially intact DNs with single molecule resolution will require further protocol developments. Third, we have only validated origamiFISH probes for targeting the common M13mp18-based scaffolds. To additionally enable origamiFISH for non-M13mp18 scaffold origami and non-origami based DNA nanostructures in the future, we include a list of probe design considerations in the Methods section. Additionally, we note that multiplexed origamiFISH imaging should be feasible with sequence barcoded DNs, which is an exciting prospect for enabling direct head-to-head comparisons between multiple DN designs in the same biological sample for lowering variability. Together, we believe origamiFISH will enable discovery of novel DN biology and design principles for eventual translation into clinically relevant nanomedicines or cellular devices.

## Methods

### DNA nanostructure synthesis

All structures were folded according to published protocols (Bastings et al., 2018), using a ThermoFisher ProFlex PCR system. ssDNA staples strands were purchased from IDT. Structures were held at 4°C following folding, until purification.

Barrel: 10 nM of p7308 ssDNA scaffold (Guild Biosciences, D441-020-1mL100) was thermo-ramped in 10x excess of ssDNA staples strands: 65°C for 15 minutes, 50 - 40°C for 98 min and 11 seconds per °C (18 hours in total). Reaction folding buffer contained 1x modified TE buffer (5mM Tris, 1mM EDTA pH8) with 10mM MgCl2.

Rod: 10 nM of p7308 ssDNA scaffold was thermo-ramped in 10x excess of ssDNA staples strands: 80°C for 5 minutes, 65 - 25°C ramp over 18 hours in total. Reaction folding buffer contained 1x modified TE buffer (5mM Tris, 1mM EDTA pH8) with 8mM MgCl2.

Rectangle: 10 nM of M13mp18 ssDNA scaffold (Bayou Biolabs, P-107) was thermo-ramped in 20x excess of ssDNA staples strands: 80°C for 5min., 60-4°C for 3.16 minutes per °C (3 hours in total). Reaction folding buffer contained 1x modified TE buffer (5 nM Tris, 1mM EDTA pH8) with 12.5mM MgCl2. Cy3-labeled rectangles were made by incubating cy3+ anti-handles (GGGATAAGTTGATTGCAGAGC/3Cy3Sp/) with folded structures for 1 hour at room temperature, following a first round of PEG purification.

### Nanostructure purification

Barrel: 10 nM of DN Barrel was incubated 1:1 (volume) with 2X PEG stock (w/w%, 10% Polyethylene glycol 8000 (Bioshop, Cat#PEG800), 1X TE buffer, 250mM NaCl, and 10mM MgCl2. Sample was incubated at room temperature for 30 mins. Sample was precipitated via centrifugation at 16,000g for 40 mins at 25°C. Supernatant was then removed with a pipette. Sample was then re-spun at 16,000g for 2 mins at 25°C, and any residual supernatant was removed with a pipette. Pelleted origami was resuspended in 1/4 volume of the original crude sample, mixed, and incubated in a thermoshaker in 1x folding buffer (Heating thermoshaker ThermoMixer® F series Model F0.5) at 30°C at 350 RPM for 20 mins to aid resuspension.

Rod: 10 nM of DN rod was incubated 1:1 (volume) with 2X PEG stock (w/w%, 10% Polyethylene glycol 8000, 1X TE buffer, 250mM NaCl, and 8 mM MgCl2. Sample was incubated at room temperature for 30 mins. Sample was precipitated via centrifugation at 16,000g for 40 mins at 25°C. Supernatant was then removed with a pipette. Sample was then re-spun at 16,000g for 2 mins at 25°C, and any residual supernatant was removed with a pipette. Pelleted origami was resuspended in 1/4 volume of the original crude sample, mixed, and incubated in a thermoshaker at 30°C at 350 RPM for 20 mins to aid resuspension.

Rectangle: 10 nM of DN rectangle was incubated 1:1 (volume) with 2X PEG stock (w/w%, 15% Polyethylene glycol 8000 (Bioshop, Cat#PEG800), 1X TE buffer, 500mM NaCl, and 12.5mM MgCl2. Sample was incubated at room temperature for 10 mins. Sample was precipitated via centrifugation at 14,000g for 30 mins at 4°C. Supernatant was then removed with a pipette. Sample was re-spun at 14,000g for 2 mins at 4°C, and any residual supernatant was removed with a pipette. Pelleted origami was resuspended in 1/4th volume of crude sample introduced to precipitation reaction, and incubated in a thermoshaker at 30°C at 350 RPM for 20 mins to aid in resuspension. Cy3-labeled rectangles were additionally purified twice using the same protocol following Cy3 labeling.

All purifications were repeated twice prior to cell uptake experiments to ensure excess staples are removed, and the final purified structures are confirmed on a 2% agarose gel to ensure purity. Concentration of the sample was obtained using a nanodrop (Thermo Scientific™ NanoDrop™ One Microvolume UV-Vis Spectrophotometer).

### Agarose gel electrophoresis

To prepare 2% agarose gel, 2.4g of agarose (FroggaBio, Cat#A87) was added to 120 mL 0.5X TBE and microwaved until dissolved. After cooling, 1.2mL of 1M MgCl2 and 5uL of Sybr-Safe (Invitrogen, S33102) were added to the solution and the gel was cast. All gels were run at 60V for 2h on ice.

### Negative stain transmission electron microscopy

DN origami samples were diluted to a final concentration of 1 nM in 1X folding buffer (1X TE buffer with 8 - 12.5 mM MgCl2 depending on structure). 3.5 µL of 1nM DN origami sample was loaded onto a plasma cleaned, carbon coated, 00 mesh Ted Pella formvar stabilized with carbon transmission electron microscopy grid (5-10 nm thickness - SFR, Cat# 01754-F). Sample was left to rest on grid for 2.5 mins, then wicked away with filter paper, washed with 1X folding buffer, and then stained for 30 seconds with 2% uranyl acetate. Samples were imaged on a Talos L120C TEM.

### Coating with oligolysine-PEG

Within 1 day of PEG purification, DNs were incubated for 30 mins at room temperature with methoxy-poly(ethylene glycol)-block-poly(L-lysine hydrochloride (mPEG5K-b-PLKC10, Alamanda polymers, Cat#050-KC010). The samples were coated at a ratio of 1:1, N:P (ratio of nitrogen in amines : phosphates in DNA). Cell uptake experiments were performed 30 minutes following oligolysine-PEG coating.

### Serum degradation assay

DNs were folded and purified as previously described and resuspended in 1X FB such that the structures had a final concentration of 50 nM. 2uL of 50nM DNs were added to 98uL DMEM + 10% FBS and incubated at 37°C for 8 hours or 2uL of 50nM was added to 98uL 1xFB for non-serum control. Samples were run on a 2% agarose gel on ice.

### PDL coverslip treatment

Coverslips were PDL functionalized up to 4 weeks prior to use. 12 mm #1.5 coverslips (Fisher Scientific cat# NC1129240) were cleaned with 1M HCl at RT overnight, and washed extensively with water to neutralize pH. Coverslips were submerged in 0.1 mg/mL PDL (Sigma; cat# P0899-50MG) overnight at RT. Coverslips were washed 5x with TC grade water, and stored in water until use at 4°C.

### Cell uptake experiments

For cellular uptake, RAW264.7 or HEK293 cells were seeded at a density of 200,000 or 100,000 cells per well, respectively, into 24 well plates containing PDL-treated #1.5, 12mm coverslips. DN samples were prepared by diluting them to their respective listed concentrations in a total volume of 200 µL DMEM complete media (DMEM supplemented with 10% FBS and 1% penn/strep). Uptake occurred at 37°C, 5% CO_2_.

### Probe design

Probe sequences used in this study are provided in **Supplementary File 1**. Probes were designed in house using the following parameters: 1) 25 nt (HCR v2.0) or 2x 25 nt (HCR v3.0) target region, 2) no more than 14 nt cross-binding to any off-targets in the mouse and human transcriptomes for each 25 nt target region, 3) Melting temperature between 45 - 65 °C, and 4) no more than 3 consecutive bases which are identical (i.e. probes which contain sequences AAAA, TTTT, CCCC, GGGG were removed). We note that similar design parameters can easily be used to generate additional origamiFISH probes towards non M13mp18-based scaffold sequences. Although probe design is challenged for shorter sequences as signal sensitivity is assisted by increased probe counts, origamiFISH is robust and can be performed with as little as 5 probe pairs (unpublished data).

### origamiFISH

Following DN uptake, cells were fixed with 4% PFA for 10 min at RT, followed by 3 × 10 min washes with 1x PBS. Samples were permeabilized with 70% ethanol either for 1 hour at RT or stored for up to one year in 70% ethanol at -20°C (sealed with parafilm to prevent evaporation).

Following permeabilization, samples were allowed to dry completely and placed, cell side down, onto 100 µL of 30% formamide hybridization buffer (molecular instruments). DN denaturation occurred at 50 °C for 10-15 minutes on a hot plate. 1 nM probe hybridization buffer was prepared in hybridization buffer. Samples were allowed to hybridize overnight at 35°C in a humidified chamber (i.e. extra wells in 24 well plate were filled with water). 3x washes were performed in wash buffer (molecular instruments) at 37°C, followed by 2x rinses in 2x SSCT (0.1% Triton-X in 2x SSC) at RT.

HCR hairpins were prepared by heating to 95°C for 90 seconds, and allowed to cool to room temperature slowly over 30 minutes to form metastable hairpins. Amplification solutions were prepared by mixing 2 µL of H1 and 2 µL of H2 hairpin in 100 µL of amplification buffer for each sample (Molecular Instruments). Washed samples were pre-amplified in 100 µL of amplification buffer for around 1 minute, and amplified in 100 µL of hairpin solution for 3 hours at RT in the dark. Following amplification, samples were washed 3 x in 2X SSC, stained with 1:3000 DAPI (ThermoFisher) for 10 minutes at RT, and mounted in Fluoromount G (Southern Biotech).

origamiFISH for lymph nodes (LNs) and brain slices were performed with the following modifications. Sample permeabilization occurred at 4°C overnight by submerging tissue sections in ice cold 70% ethanol as free-floating sections in a 24 well plate (or in a coplin jar for LN cryosections on slides). Following permeabilization, sections were rinsed 3x in 1x PBS to remove traces of ethanol. This is important as ethanol precipitates with the origamiFISH hybridization buffer. DN denaturation occurred at 60°C for 15 minutes on a hot plate, followed by hybridization in 1nM probe solution at 35°C in a humidified chamber. HCR hairpin amplification occurred at RT overnight on a rocker. Vibratome sections were mounted onto slides using a paint brush.

origamiFISH for whole-mount organoids were performed with the following modifications, adapted from previously published EASI-FISH (Wang et al., 2021b) and CLARITY protocols (Shah et al., 2016). Whole organoids were permeabilized with 70% ethanol for at least 2 days in a 96 well-plate at 4°C. Following permeabilization, organoids were subjected to DN denaturation in hybridization buffer at 60°C for 30 minutes. Following 3x rinses in 1x PBS, organoids were incubated in MOPS buffer (Bioshop, 20mM, pH 7.7) for 30 minutes. DNs and endogenous nucleic acids were covalently tethered to MelphaX by incubating organoids overnight at 37°C in MOPS buffer supplemented with 1 mg/L MelphaX and 0.1 mg/mL Acryloyl-X, SE (100 µL). MelphaX and Acryloyl-X SE were prepared as previously described (Wang et al., 2021b). The next day, organoids were rinsed in 1x PBS.

4% acrylamide solution was prepared for sample clearing containing 4% (v/v) 19:1 acrylamide/bis-acrylamide, 60nM Tris-HCl pH 8, and 0.3M NaCl. The acrylamide solution was degassed for 10 minutes on ice using nitrogen gas. Organoids were acclimated to the acrylamide solution for 15 minutes on ice. 10% (w/v) ammonium persulfate (APS) was freshly prepared and kept on ice until use. A 4% hydrogel solution was prepared by mixing the 4% acrylamide solution with 0.05% APS and 0.15% TEMED. Organoids were introduced to 4% hydrogel solution and immediately pulled into a lightsheet capillary for gellation. Capillaries containing the organoid/gel solution were allowed to polymerize for 2 hours at 37°C in a humidified incubator.

Clearing buffer was prepared by mixing 1:100 proteinase K, 50mM Tris-HCl pH 8, 1mM EDTA, 0.5% Triton-X, 500nM NaCl, and 2% SDS. Following gellation, organoid-gels were carefully trimmed and washed in 2X SSCT (0.5% Triton-X). SSCT was then replaced by the clearing buffer, and samples were allowed to clear for 3 days 37°C with rocking (350 RPM). After clearing, samples were washed 3 x in 2X SSCT prior to probe hybridization and amplification.

For hybridization, organoid-gels were first equilibrated in hybridization buffer for 30 minutes at 35°C, followed by overnight hybridization at 35°C in 1nM probe solution in hybridization buffer. The next day, organoid-gels were washed in wash buffer 3x 20 minutes at 37°C, followed by 2x 15 minute washes with 2x SSCT. Amplification occurred at RT overnight, followed by 3x 20 minute washes with 2x SSCT, and staining in 1:3000 DAPI solution for 3 hours. Organoid-gels were taken directly to lightsheet imaging.

### Cellular transfections

HEK293 cells were transfected with lipofectamine 3000, and RAW264.7 cells were transfected with Fugene 6, following publisher’s protocols. All plasmids were introduced at 1:3 ratio.

### Immunostaining

Cells: Immunostaining was performed following published protocols (Wang and Lefebvre, 2022). Briefly, cells were blocked in blocking buffer (4% NDS in 0.1% PBST) for 1 hour at RT. Primary incubation occured at 4°C overnight. Following 3x 15 minute 0.1% PBST washes, samples were incubated for 3 hours at RT with alexa-conjugated secondary antibodies (Invitrogen or Jackson ImmunoResearch). Samples were mounted onto glass slides using Fluoromount G (Southern Biotech).

Tissue: Immunostaining of tissue slices were performed following protocol above, with a few modifications. Primary incubation occurred at 4°C for 3 nights. Samples were incubated in secondary antibodies for 3.5 - 4 hours at RT. origamiFISH was performed immediately following immunostaining. Antibodies and concentrations used in this study were as follows: 1:1000 rat anti-LAMP1 (DSHB, 1D4B); 1:1000 rabbit anti-PHH3 (Millipore Sigma, cat# 06-570); 1:400 rat anti-F4/80 (ThermoFisher, cat# 14-4801-82). Nuclei were labeled using DAPI at 1:3000 in 2x SSCT for 10 minutes at RT.

### Turbo origamiFISH

The standard origamiFISH protocol was followed, with the following modifications: 1) cells were fixed and permeabilized in 100% methanol for 10 min at -20°C, 2) Probe hybridization occurred at 1 µM for 5 minutes, 3) all washes were shortened to 5 min each, 4) HCR amplification occurred for 30 min at RT.

### Confocal microscopy

Images were acquired on a Leica SP8 scanning confocal microscope, using a 40X oil objective (NA = 1.3), using a tunable white light laser. Z-stacks were collected with a 0.5 µm step size throughout the depth of the cells. RAW264.7 cells were imaged using 1.28x magnification. HEK293 cells were imaged using 1x magnification. Laser power and gain used were: Alexa 488: 12% / 30%, Cy3/Alexa 568: 12% / 30%, Alexa 647: 10% / 50%. Images of tissue slices were acquired with optimized laser powers using the Z-compensation feature throughout the depth of the tissue. All images were either acquired with the Leica Lightening module for integrated image deconvolution at the time of imaging, or deconvolved using Huygens deconvolution.

### Lightsheet microscopy

2% low melting point (LMP) agarose was made in water, and stored at RT until use. Cerebral organoids were cleared and mounted in 2% LMP agarose. The agarose solution containing organoid was pulled into a glass capillary tube, allowed to cool, and submerged into imaging solution in the imaging chamber on a Zeiss Lightsheet Z1 microscope. The agarose-embedded organoid was plunged out of the capillary tube directly into the imaging buffer. Images were acquired using a 20X (NA = 1.0) objective with dual-side illumination, with 0.8µm step sizes throughout the depth of the organoid. Images were merged and processed in Bitplane Imaris.

### Mouse strains

Wild-type C57/B6J mice used in this study were purchased from Jackson Laboratories or Charles River Laboratories. All experiments were carried out in accordance with the Canadian Council on Animal Care guidelines for use of animals in research and laboratory animal care under protocols approved by the Laboratory Animal Services Animal Care Committee at the Hospital for Sick Children (Toronto, Canada) and the Division of Comparative Medicine (Toronto, Canada).

### Live brain slice uptake

Mice were sacrificed and mouse brains were quickly removed and immersed in ice-cold slicing solution saturated with 95% O_2_ and 5% CO_2_ containing: 75mM sucrose, 120mM NaCl, 2.5mM KCl, 1.3mM MgSO4, 1mM NaH2PO4, 26mM NaHCO3, 1.25mM CaCl2, and 11mM D-glucose.

350 µm sagittal slices were obtained using a vibratome (VT1200s, Leica). Slices were recovered at 32°C for 45 min to 1 hour and then stored at RT, in artificial cerebral spinal fluid (aCSF) saturated with 95% O_2_ and 5% CO_2_ containing: 120mM NaCl, 3.0mM KCl, 1.2mM MgSO_4_, 1.0mM NaH_2_PO_4_, 26mM NaHCO_3_, 2.0mM CaCl_2_, and 11mM D-glucose. Slices were transferred to a 6 well plate, in aCSF, for uptake in 1nM DN barrels, while maintaining constant bubbling with 95% O_2_ and 5% CO_2_. Slices were fixed in 4% PFA post-uptake overnight at 4°C prior to origamiFISH.

### Cortical organoid generation and DN uptake

Human iPSC line PGPC-17/11 (Personal Genome Project Canada) was cultured on Matrigel-coated plates in mTeSR Plus medium (STEMCELL Technologies 05825) and passaged with ReLeSR (STEMCELL Technologies 05873) every 5-7 days. Cells were routinely tested for mycoplasma negativity.

Cortical organoids were generated from the PGPC-17/11 hiPSC line as previously described (Pa$ca et al., 2015), with modifications. Briefly, hiPSCs were dissociated into single cells using Accutase (STEMCELL technologies 07920). On Day 0, 10,000 cells were aggregated in each well of a PrimeSurface V-bottom 96-well Plate (S-Bio) to form embryoid bodies (EBs), in mTeSR Plus medium containing ROCK inhibitor Y27632 (10uM), for the first 24 hours. From Day 1-6, EBs were fed with Essential-6 medium (Thermo Fisher Scientific A1516401) containing 2.5uM dorsomorphin (Tocris) and 10uM SB-431542 (Tocris). From Days 6-25, medium was changed to NGD medium (Neuro-glial differentiation medium) supplemented with 20ng/mL EGF (Peprotech) and 20ng/mL FGF2 (Thermo). From Days 25-43, medium was changed to NGD medium supplemented with 20ng/mL. BDNF (Peprotech) and 20ng/mL NT-3 (Peprotech). From Day 43 onwards, organoids were grown in only NGD medium.

NGD was prepared as previously described (Li et al., 2019). 500 mL of 0.5x NGD consists of 475 mL of Neurobasal (Thermo), 5mL of Gem21-VitA (0.5X, Gemini Bioproducts), 2.5mL of Neuroplex N2 (0.5X, Gemini Bioproducts), 5mL of 100mM pyruvate, 5mL of Glutamax, 5mL of Pen/Strep, 5 mL of 5M of NaCl, 1g of Albumax I, 3.5µg of Biotin, 85mg of Lactic acid, and 2.5mg of Ascorbic Acid.

12 week old organoids were subjected to DN uptake through introduction of 1nM DN barrels in 200µL of NGD. Uptake occurred at 37°C under 5% CO_2_. Following uptake, DNs were post-fixed in 4% PBS for 2 hours at RT, followed by origamiFISH.

### Lymph node (LN) injection and collection

DN barrels were introduced intramuscularly following published protocols (Rezende et al., 2019). Four hours following uptake, the inguinal lymph nodes were dissected and fixed overnight in 4% PFA overnight at 4°C. Fixed LNs are washed in 1x PBS 3 times for 10 minutes each. LNs were sectioned on a vibratome (Leica) at 100 µm, prior to origamiFISH and immunostaining as free-floating sections in a 24 well plate.

### Image analysis

Image processing and analyses were performed in Bitplane Imaris (v 9.9.0) and FIJI. Data analysis and visualization were performed in Graphpad Prism (version 9) or using the Seaborn library in python.

Images were preprocessed in FIJI by performing the following operations to generate an additional, synthetic autofluorescence channel for cell segmentation: 1) origamiFISH channel was gaussian blurred with sigma = 4; 2) origamiFISH channel was median filtered with radius = 7; 3) image calculator was used to add the origamiFISH and DAPI channels; 4) Added channel was gaussian blurred with sigma = 4.

Images were imported into Bitplane Imaris for downstream analyses using the cell module for single cell spot counts and differential localization to the cytoplasm and nucleus, and the surfaces module for spot volume analyses. Parameters were optimized for each experiment and maintained for all samples within the experiment.

## Acknowledgements

W.X.W acknowledges the PRiME initiative for fellowship support. T.R.D acknowledges the Barbara and Frank Milligan Graduate fellowship. Y.L acknowledges NSERC Discovery Grant, Medicine by Design, Simons Foundation, Brain Canada, Stem Cell Network, Can-GARD, and Brain & Behavior Research Foundation. J.M. acknowledges NSERC Discovery Grant and The Hospital for Sick Children. L.Y.T.C acknowledges NSERC Discovery Grant, Medicine by Design, and the Canadian Foundation for Innovation for funding. We thank members of the Chou lab for helpful comments on this manuscript. We thank Paul Paroutis and the Hospital for Sick Children imaging facility for instrument support and guidance on lightsheet microscopy. We thank Dr. Julie Lefebvre for the generous gift of antibodies, reagents, and use of equipment.

## Author Contributions

W.X.W and L.Y.T.C conceptualized the project and designed the experiments. W.X.W performed experiments and computational analyses with assistance from T.R.D, H.Z, A.B and M.R. Z.P.J, J.M, Y.L and L.Y.T.C participated in data interpretation and provided supervision. W.X.W and L.Y.T.C wrote the manuscript with input from all authors.

---

**Supplementary Fig. 1.**
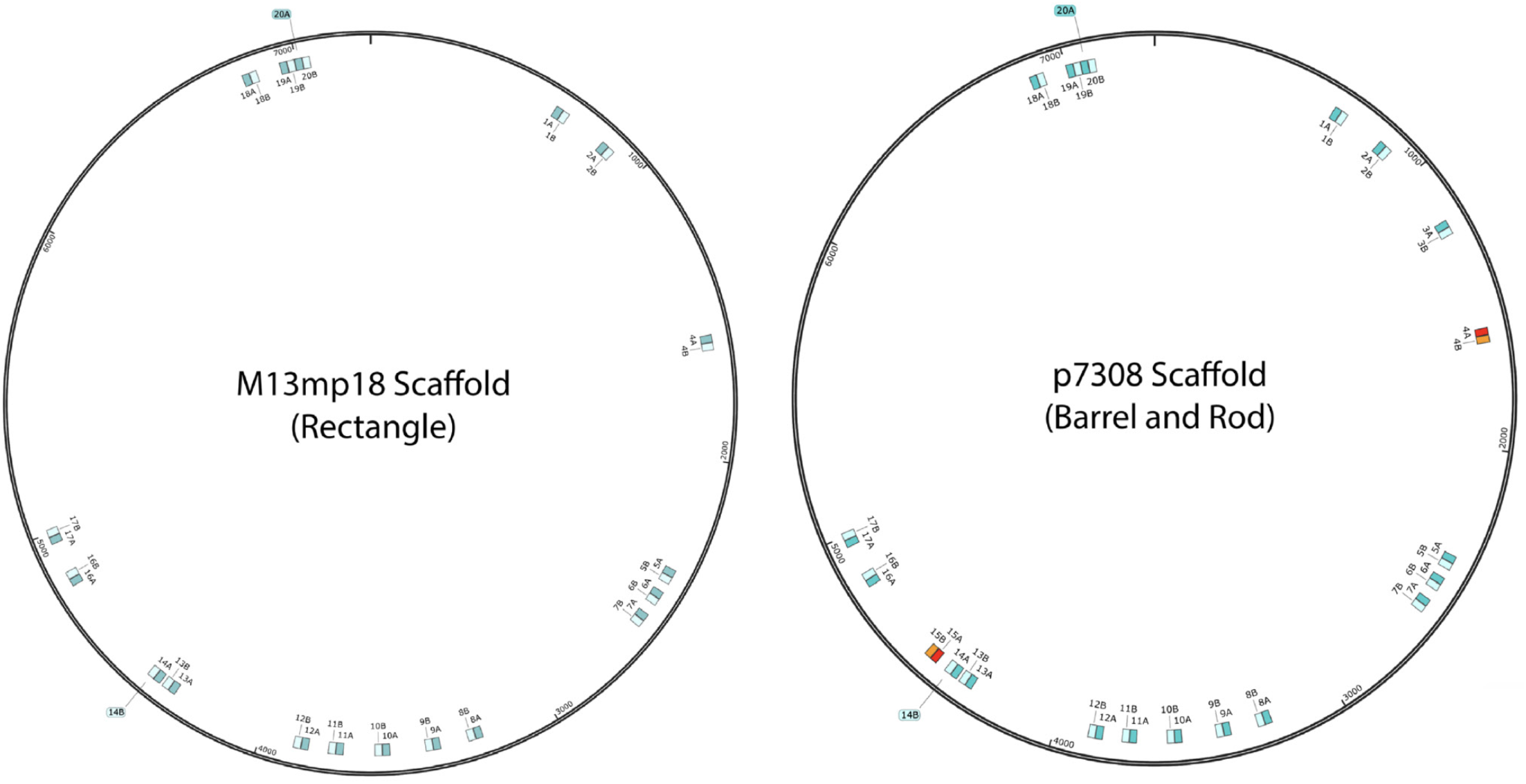
Probe design overview. 18 split-initiator probe pairs bind to the M13mp18 scaffold used to generate DN rectangles (*left*). The first half of the probe pair (A) is highlighted in cyan. The second half of the probe pair (B) is highlighted in light blue. Binding sites for probes 4 and 15 are not present within the M13mp18 scaffold. 20 Split initiator probe pairs bind to the p7308 scaffold used to generate DN barrels and long rods (right). Probes 4 and 15 are unique to the p7308 scaffold and correspondingly highlighted in red/orange.

**Supplementary Fig. 2.**
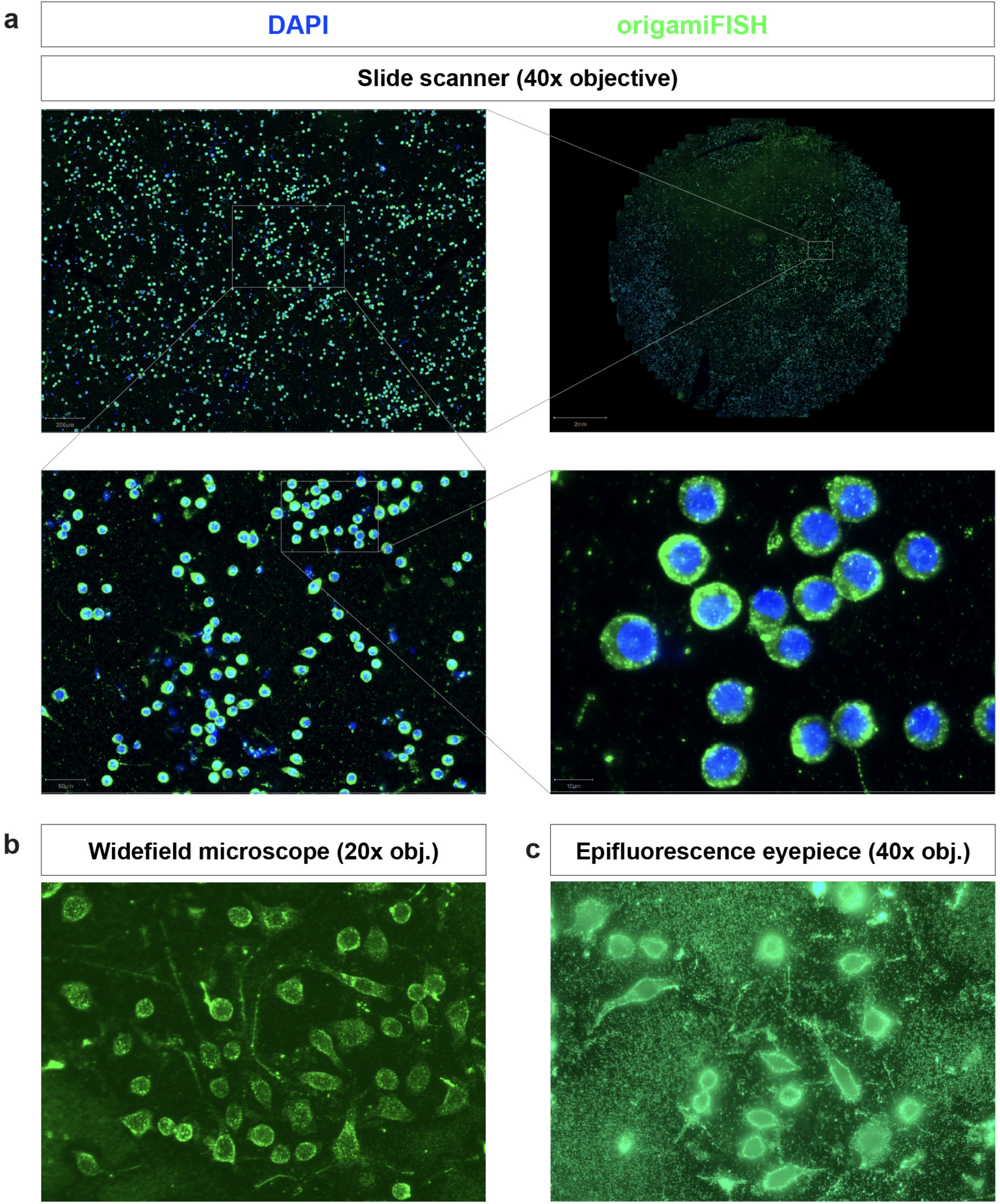
origamiFISH performs robustly across luorescence imaging platforms. **a** automated slide scanner (40x oil objective) enables large-scale, high-throughput imaging of origamiFISH staining. **b** widefield fluorescence microscope (Echo Revolve; 20x air objective). **c** Image obtained on epifluorescence eyepiece phone mount (40x oil objective). extracellular punctate staining represents negatively-charged DNs localized to positively-charged PDL treated coverslips, which are removed through confocality on confocal setups.

**Supplementary Fig. 3.**
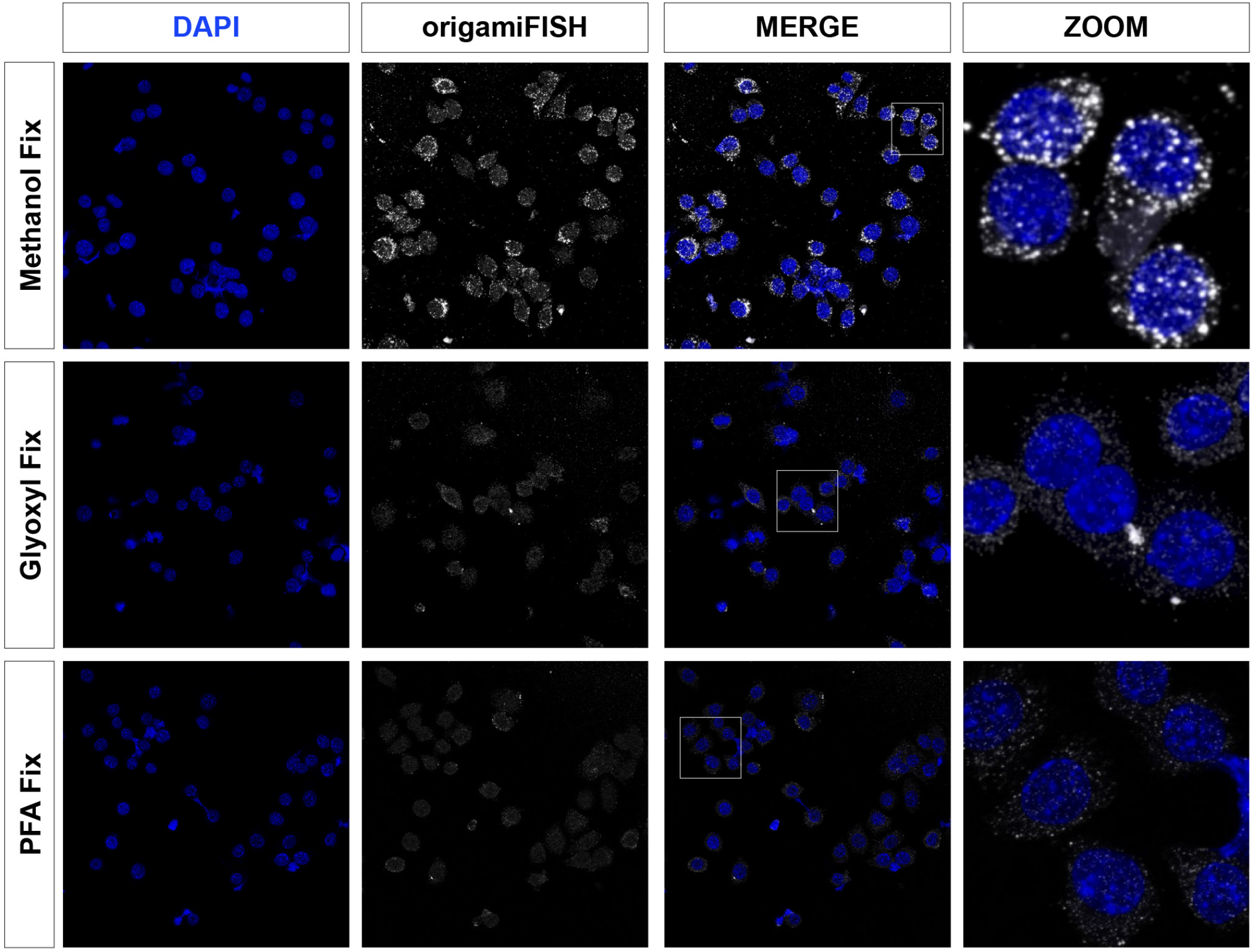
Turbo origamiFISH is dependent on methanol fixation and allows for rapid prototyping of DNs. Turbo origamiFISH shortens 1 day origamiFISH protocol to 1 hour (end-to-end). Turbo origamiFISH is dependent on methanol fixation, similar to RNA smFISH (Shaffer et al., 2013). Glyoxyl fixation (Richter et al., EMBO; 2018) was included as an alternative test to PFA fixation, but which also did not result in positive Turbo origamiFISH staining.

**Supplementary Fig. 4.**
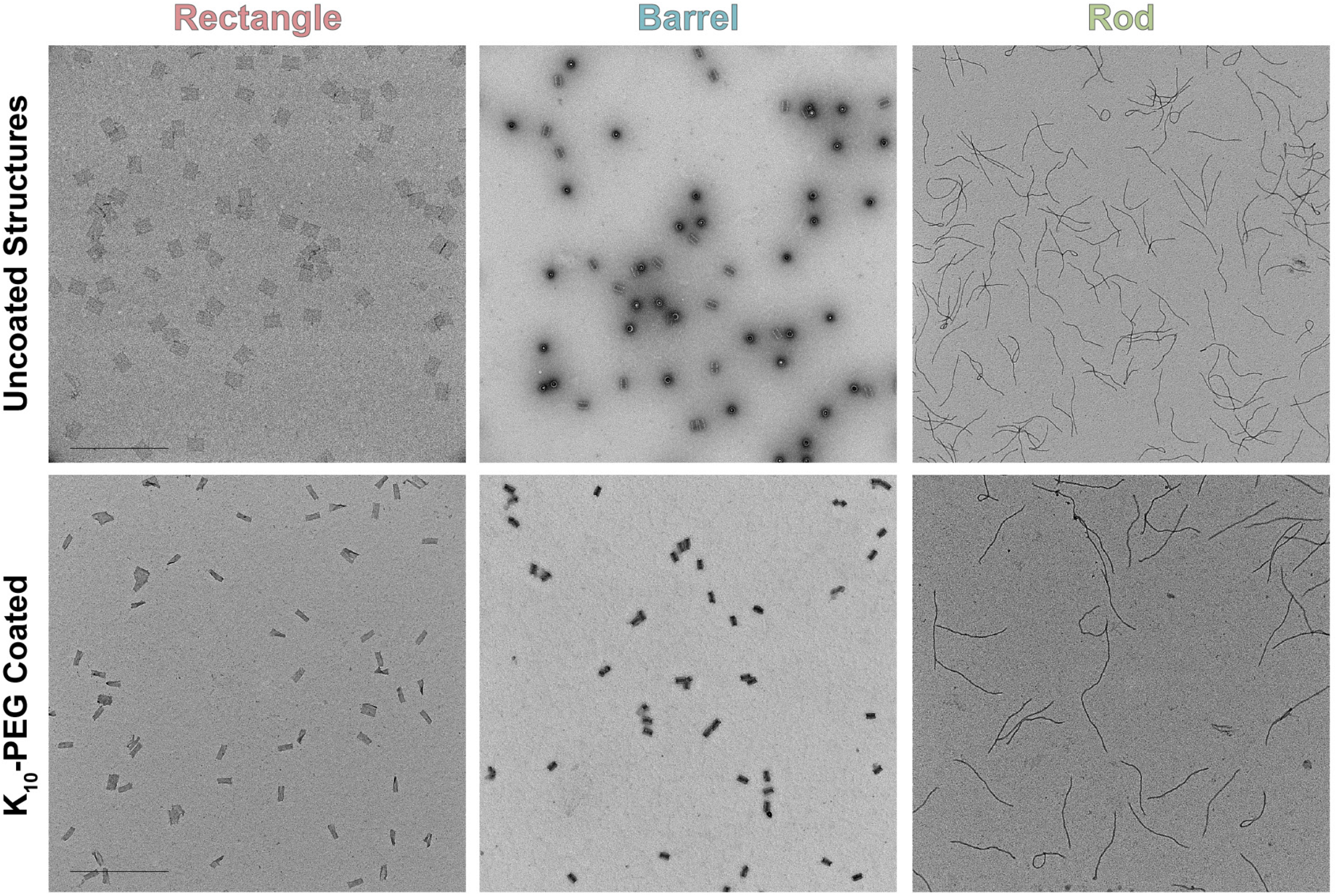
Negative stain TEM images of three DN shapes used in this study. Uncoated DNs are shown in the top panel. DNs retain shape-specific characteristics following K_10_-PEG coating (*bottom*). Scale bars are 500 nm.

**Supplementary Fig. 5.**
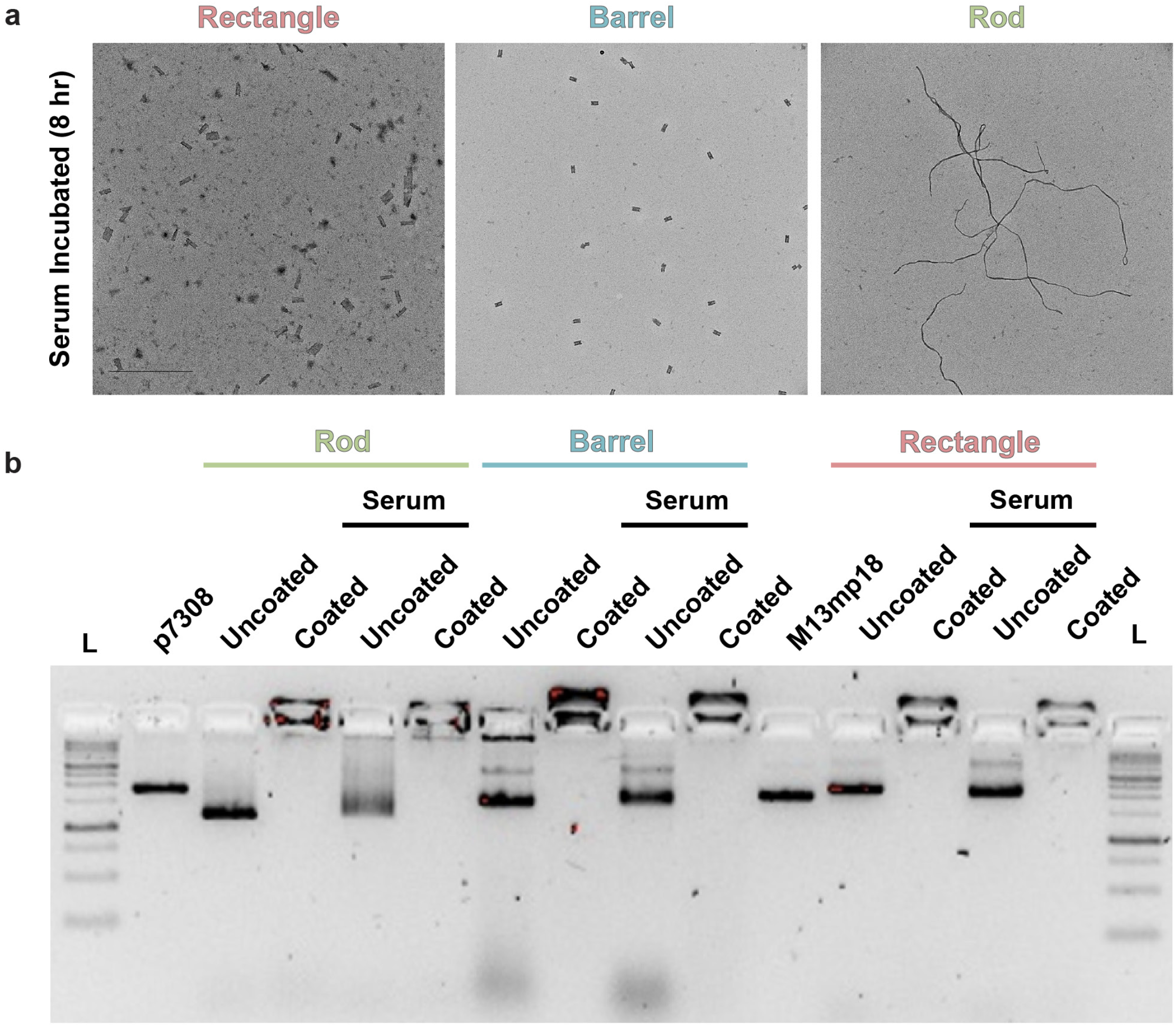
Characterization of DN integrity under cellular conditions. **a** Negative stain TEM images of K10-PEG coated structures following 8 hour incubation under serum conditions (DMEM + 10% FBS). Scale bars are 500 nm. Most structures retained their shape following serum incubation, although a subset of DN rectangles appear to have degraded. **b** Effect of K10-PEG coating and serum incubation (8 hours) on DNs, as assessed through agarose gel electrophoresis. Coated: 1:1 K_10_-PEG coated structures; serum: serum incubated over 8 hours; L: ladder; p7308: p7308 scaffold alone; M13mp18: M13mp18 scaffold alone. The major DN band is retained for all DN structures following serum incubation. Gels were run for 2 hours on ice.

**Supplementary Fig. 6.**
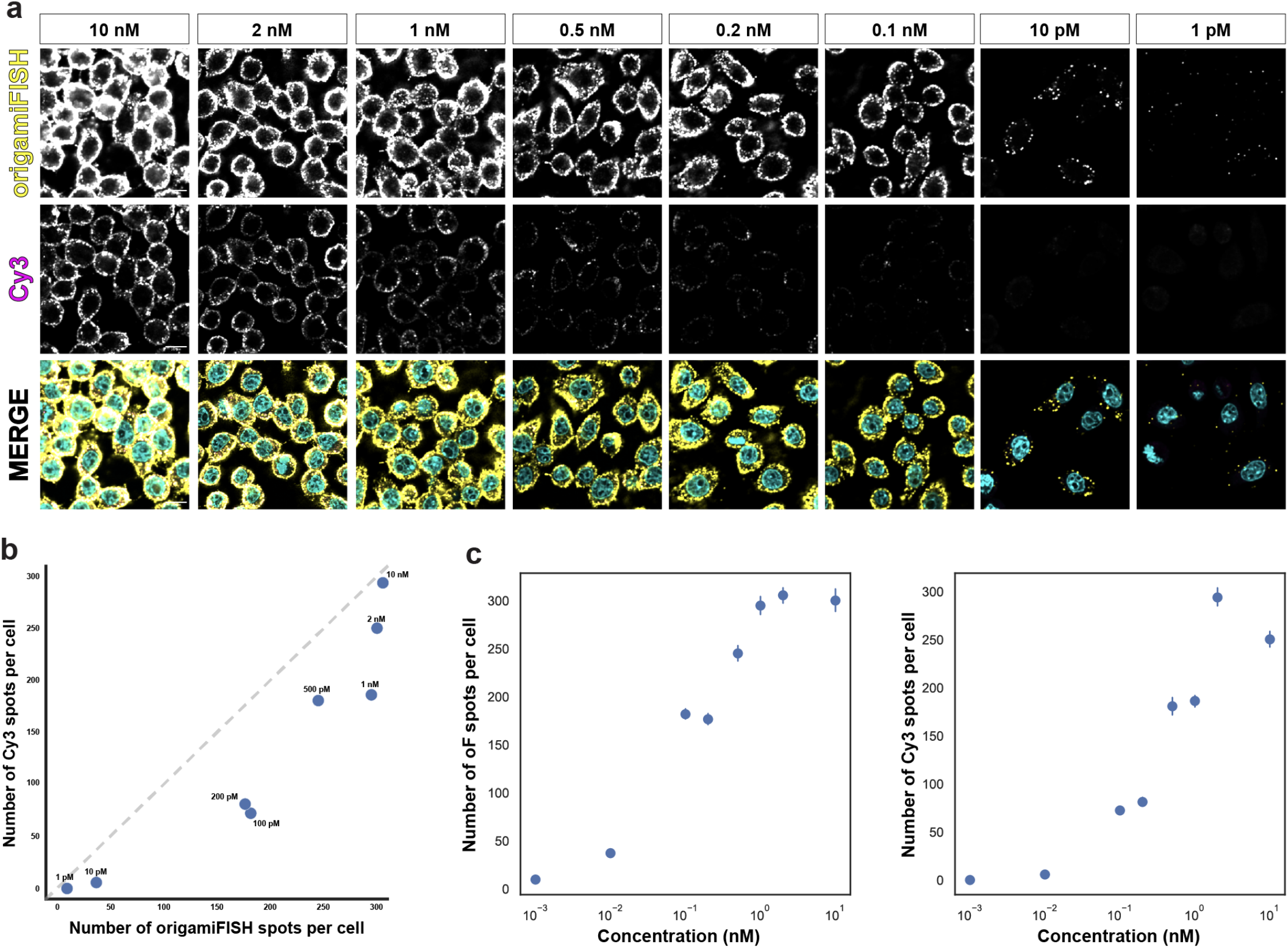
origamiFISH is more sensitive compared to dye labels across concentration gradient. **a** Representative images for origamiFISH (yellow) and Cy3 (magenta) across all concentrations analyzed. **b** Correlation between the number of origamiFISH spots per cell (x) and the number of Cy3 spots per cell (y). Dashed line represent data distribution for equal sensitivity between origamiFISH and Cy3. Data points which fall below the line indicate higher origamiFISH sensitivity. Data points which fall above the line indicate higher Cy3 sensitivity. origamiFISH demonstrate superior sensitivity at all concentrations. **c** the number of origamiFISH (left) and Cy3 (right) spots plotted against uptake concentration. Both demonstrate a dose response relationship between uptake concentration and spot number. Signal saturation is observed past the 2nM concentration for both origamiFISH and Cy3.

**Supplementary Fig. 7.**
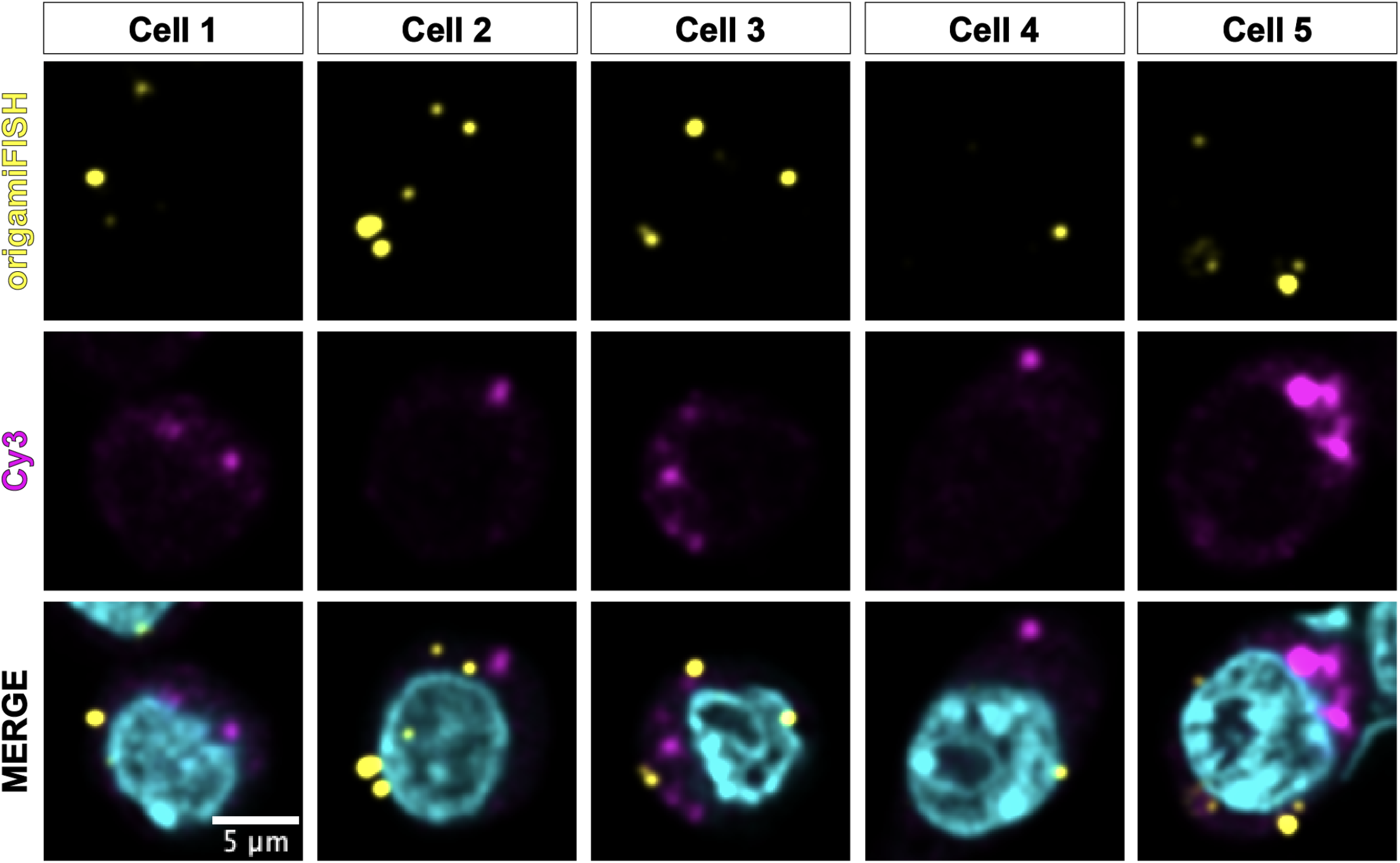
origamiFISH and Cy3 dye labels do not colocalize following low concentration DN uptake. Additional example images of Cy3+/origamiFISH- and origamiFISH+/Cy3-spots following 1pM DN uptake.

**Supplementary Fig. 8.**
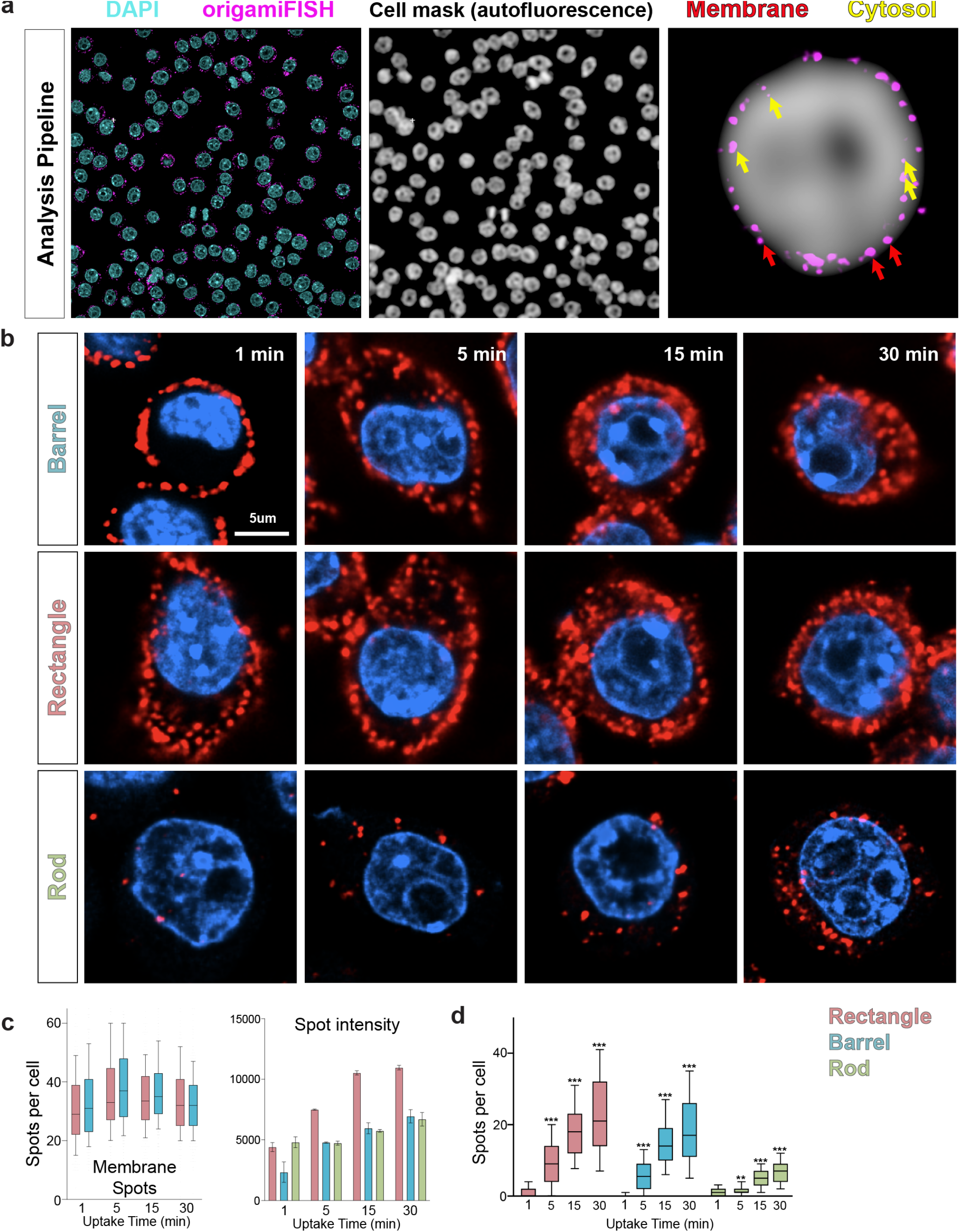
Summary of DN internalization dynamics and analyses pipeline. **a** Cell masks were created using the combined cellular autofluorescence from the origamiFISH and DAPI channels. Signal addition and blurring was used to create a cellular mask channel, which was utilized to generate cell membrane surfaces for analyses. Membrane spots were characterized as spots which fall along the boundary of the cell mask channel. Cytosolic spots were characterized as spots which fall inside the cell mask. **b** Representative images for three shapes at four uptake time points. **c** left: Number of membrane localized spots quantified for DN rectangles and barrels are relatively consistent across the time-course, suggesting early adsorption onto the cell membrane. *right*: origamiFISH spot intensity increases over the uptake timecourse for all three DN structures, as expected. **d** The number of spots per cell increases with uptake for all three structures. *p<0.05, **p<0.005, ***p<0.00001; comparisons are between the sample with the previous time-point (i.e. 5 minute barrel uptake is compared to 1 minute barrel uptake; 15 minute barrel uptake is compared to 5 minute barrel uptake).

## Notes

### Competing Interest Statement

The authors have declared no competing interest.

### Summary of Updates

Correct page sizes to remove watermark which was overlayed on top of figures.

